# Spatiotemporal cell type deconvolution leveraging tissue structure

**DOI:** 10.64898/2026.02.10.705204

**Authors:** Macrina Lobo, Ziqi Zhang, Xiuwei Zhang

**Author notes:** Contributing authors.

## Abstract

Spot-based spatial transcriptomics (ST) captures aggregated transcriptomic profiles at spatial locations (spots) in tissue slices. Cell type deconvolution methods decode each spot and estimate the proportion of every cell type in the spot, necessary for uncovering spatial cell type distributions for further downstream analyses. Existing methods utilize cell type markers or reference transcriptomic (scRNA-seq) atlases at single cell (sc) resolution, or by aggregating profiles of identified cell types. However, current methods fail to effectively utilize the 3D tissue layout and single cell resolution reference. Some leverage 2D spatial organization assuming proximal spots are similar, which may be violated around boundaries or isolated cell types. We present SpaDecoder, a parallelized matrix factorization-based per-spot deconvolution method for multiple 3D spatial or temporal ST tissue slices effectively leveraging tissue structure with an adaptively inferred 3D neighborhood Gaussian kernel. We additionally account for variability in sc-reference profiles, along with batch effects.

The mathematical framework of SpaDecoder allows it to be used for a range of downstream analyses. It can decode anteroposterior variability, impute gene expression, uncover putatively key tissue regions, identify colocalized cell types and predict spatio-temporal scRNA-seq cell locations. Ablation tests along with comparisons against other methods on various metrics, datasets, and scenarios, collectively show that SpaDecoder effectively harnesses 3D tissue structure and sc-reference profiles to improve cell type deconvolution. SpaDecoder is available at https://github.com/ZhangLabGT/spadecoder.

## Introduction

Higher order multicellular organisms typically comprise trillions of single cells that are spatially organized into complex tissues with diverse structures performing specific functions. Each cell contains millions of RNA molecules which, when measured via single cell RNA sequencing (scRNA-seq) and mapped to genes, provide insights into the gene expression driving cellular state and function, cell-type markers, and identification of novel cell types not evident from tissue morphology. Capturing RNA molecules along with their spatial locations further elucidates the 3D organization of cells into tissues, the spatial expression pattern of genes, as well as intracellular and intercellular communication signaling pathways[1]. These insights are critical to several biological questions such as the underlying mechanisms driving development and healthy cellular processes as well as disruptions due to diseases such as cancer. Additionally, developmental and stem cell biologists may utilize ST to inform *in vitro* stem cell differentiation efforts to 3D organoid models to study developmental disorders[2].

Several spatial transcriptomic (ST) technologies have been developed to measure RNA levels along with spatial location at spot, cellular, or sub-cellular resolution by capturing individual RNA molecules [3]. ST technologies typically offer trade-offs between cost, number of genes profiled, spot resolution and the detection efficiency of genes. A single spot may contain 10-200 cells for older ST technologies or 1-30 cells for newer technologies such as Visium from 10x Genomics, or may be at subcellular resolution [4, 5]. Compared to single cell or subcellular spatial omics, multi-cell spots typically captures more genes, at higher detection efficiency and lower cost and is also most abundantly available. However, to utilize this, computational techniques are needed to decode the spots into the individual cell-types.

Another different but sometimes complementary computational task is determining the spatial coordinates of cells from an scRNA-seq dataset. Deconvolution methods utilizing scRNA-seq reference atlases sometimes perform mapping of cells in the reference with cells or spots in the ST dataset as a part of the algorithm [6]. With such a mapping, it becomes possible to estimate the location of scRNA-seq cells and predict the expression of genes that may not be spatially captured in the ST data. Further, denoising can be performed on spatially expressed genes.

Methodologically, existing spatial cell type deconvolution methods can be categorized into probabilistic [7– 13], NMF-based [14–16], graph-based [17, 18], deep-learning [6] or Optimal Transport based[19, 20] with some overlap between categories [21]. Current approaches for spatial deconvolution utilize scRNA-seq reference atlases [21–24] which provide single cell resolution transcriptome wide profiles of the same tissue region or are reference free [25–27]. Some reference-based approaches have reference-free modes of operation [11]. Since reference based methods with appropriately chosen reference datasets typically outperform reference-free methods [11], it behooves us to utilize abundantly available scRNA-seq atlases while handling the additional complexity arising out of batch effects between scRNA-seq and spatial datasets due to different sequencing depth, gene detection efficiency, and dropout. The scRNA-seq atlas should capture a similar section of tissue at the same developmental, healthy or diseased state to prove most effective.

While a number of computational methods have been developed, existing methods have several limitations. First, among reference-based methods with the exception of a few like the single-cell version of tangram [6], most approaches aggregate single-cell profiles across cells of the same type thereby failing to fully utilize the diversity in cell state that arises even among cells from a predefined cell-type. Incorporating reference scRNA-seq at single cell resolution would help in this regard. Second, although an increasing number of experiments profile multiple spatially proximal tissue slices in order to study 3D cellular and molecular behavior within their spatial context, there is no method that takes advantage of multiple spatial samples that are adjacent in a tissue or temporally related. Third, while recent work [11, 28] has leveraged the 2D spatial proximity of spots, under the hypothesis that spatially adjacent spots have similar cell composition, this assumption while true in more homogeneous cellular environments, can face limitations with more complex tissue structures[29]. It is therefore useful to share information across tissue locations that are spatially as well as transcriptionally similar while adapting to the heterogeneity of the region surrounding each spot.

In order to address the aforementioned challenges, in this work we present SpaDecoder, which predicts cell-type proportions by optimizing a matrix factorization based loss function. SpaDecoder uses a 3D weighted spatial Gaussian kernel which enables information sharing across multiple adjacent slices as well as within the same slice. The kernel is adaptive which allows for the selection of transcriptionally similar neighboring spots, and alleviates the problem of dissimilar neighboring spots worsening the deconvolution performance. Specifically, in order to adapt to both homogeneous as well as heterogeneous tissue environments, we develop a permutation based localized weighted spatial autocorrelation metric which when applied to each spot intelligently selects the neighborhood within which to pool transcriptomic and spatial information to improve deconvolution predictions. Finally, SpaDecoder leverages individual single cell reference profiles (as opposed to cell-type aggregated) from the scRNA-seq atlas.

We first establish the deconvolution performance of SpaDecoder qualitatively as well as quantitatively under various scenarios using three spatial datasets. SpaDecoder consistently outperforms baseline methods in all scenarios. Next, we define several metrics and perform downstream analyses to distill SpaDecoder cell type proportions and scRNA-seq-spatial maps into interpretable biological findings. We showcase the application of SpaDecoder on several downstream tasks. We capture antero-posterior cell type variability across 12 samples or slices of the mouse hypothalamic preoptic region, denoise spatial gene expression and predict the expression of unmeasured genes. In human breast cancer tissue, we identify colocalized cell types and subtypes in 3D spatial tissue neighborhoods and isolate key tissue regions associated with the corresponding cell types. Finally, we demonstrate the performance of SpaDecoder on a temporal dataset capturing 16-19 post conception weeks (pcw) of human embryonic thymic development, where, in addition to identifying cell type regions and colocalized celltypes, we predict the 3D spatio-temporal locations of scRNA-seq reference cells.

## Results

### Overview of SpaDecoder

SpaDecoder is described in Methods, with technical details in Supplementary Notes and a schematic in Figure 1. It is developed for experiments profiling a 3D stack of multiple spatial or temporal spot-based spatial transcriptomic (ST) slices in proximity to each other (Fig. 1a). The input comprises “query” ST spot resolution gene expression and coordinate matrices (Fig. 1b) with a “reference” cell type annotated single cell RNA-sequencing (scRNA-seq) cell-by-gene expression matrix (Fig. 1c). If the scRNA-seq cells are not annotated, one can follow standard processing pipelines such Luecken *et al* [30] in scanpy [31] or Seurat [32] for pre-processing and clustering at a biologically meaningful resolution. Cell type annotations are one-hot encoded into a cell-by-cell-type matrix. The objective of SpaDecoder is to identify the proportion of each of the pre-defined scRNA-seq reference cell-types or clusters at each query spatial spot location. For each spot, SpaDecoder additionally outputs a reference cell-to-cell-type association matrix, that represents the probabilities of scRNA-seq cells lying on a continuum of discrete cell states; and a spot correction scalar, that represents spot specific batch effects. (Fig. 1d).

**Fig. 1.**
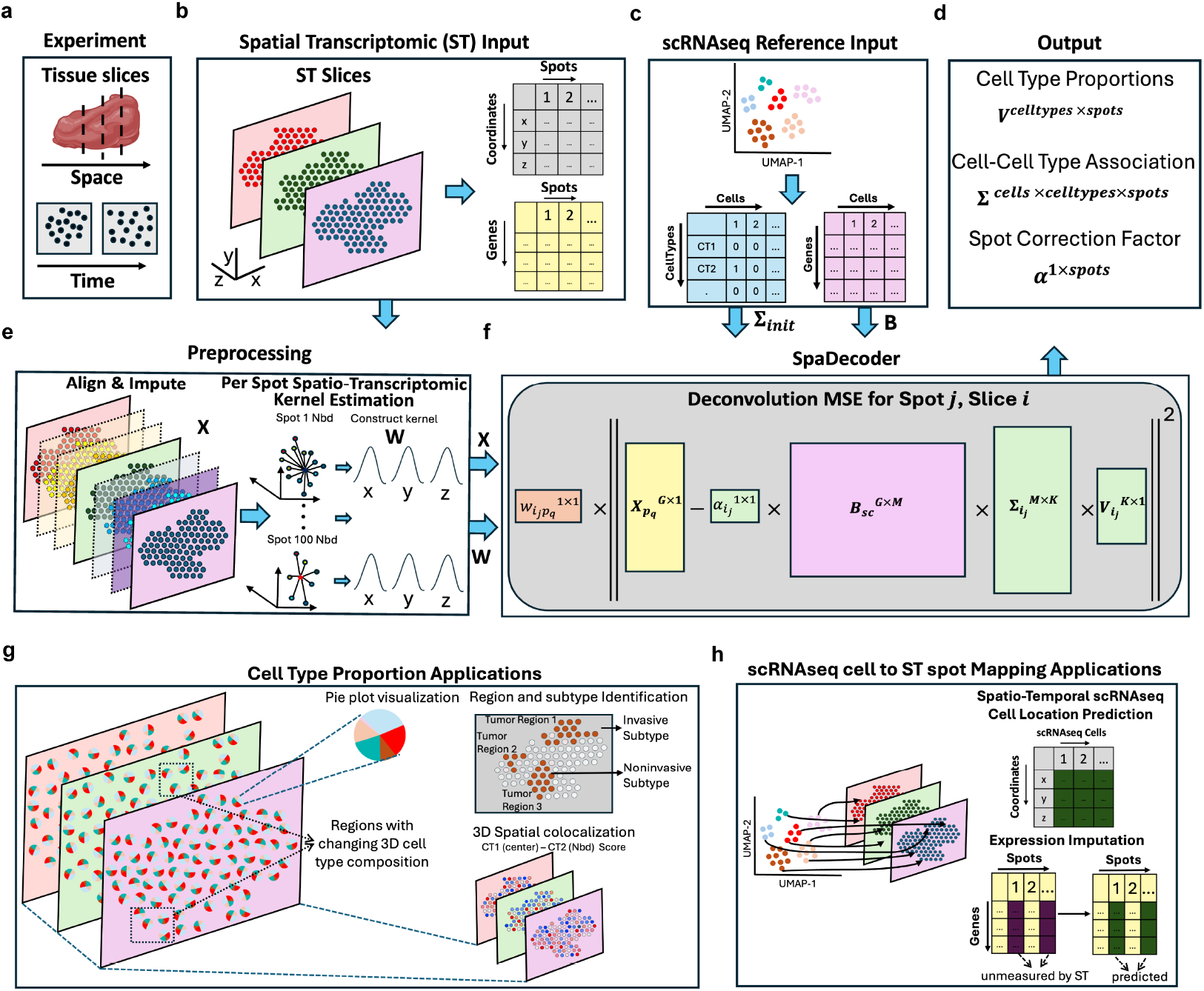
SpaDecoder performs spatio-temporal deconvolution of 3D tissue slices. Schematic of the SpaDecoder approach (a) It handles experimental data from multiple tissue slices profiled spatially or temporally (b) The spatial transcriptomic input “query” comprises a spot by (*x, y, z*) coordinate matrix and a spot by gene expression matrix (c) The scRNA-seq reference dataset is used to obtain a cell by gene expression matrix. scRNA-seq celltype annotations are used to obtain a type one-hot encoded cell by celltype matrix with 1 indicating that the cell belongs to the corresponding cell type and 0 otherwise. (d) For each spot, SpaDecoder outputs celltype proportions which sum to 1, a cell-cell type association matrix whose columns sum to 1, and a spot correction scalar. (e) SpaDecoder preprocessing comprises alignment of multiple slices with moscot [33], imputation of additional intermediate slices, per spot 3D neighborhood selection and spatio-transcriptomic kernel estimation. (f) SpaDecoder objective function for spot *j* in slice *i* (g) SpaDecoder cell type proportions for each spot are used in spot-level pie chart visualizations, identifying regions with changing cell-type composition across slices, pinpointing the spatial extent of each cell-type containing region, identifying global and local 3D spatial cell type colocalization (h) SpaDecoder infers an scRNA-seq cell to spot mapping matrix which is used to predict the spatio-temporal coordinates of single cells, to denoise existing spatial gene expression and predict expression of unmeasured genes

Our pre-processing pipeline (Fig. 1e) comprises slice alignment and intermediate slice imputation to bridge the gap between slices and augment our dataset; adaptive neighborhood inference to account for homogeneous and heterogeneous tissue regions; and spot-specific 3D Gaussian kernelization of computed spatial distances (Methods). While the spatial similarity in the 3D neighborhood around the current spot can be used to improve deconvolution of the current spot, the similarity often decays with distance[29]. Therefore, like others [11], we model this with a Gaussian kernel applied on the spatial distance between pairs of spots. While previous work [11][28] used a fixed spatial neighborhood and bandwidth, we automatically infer a set of transcriptionally similar spots that are in close spatial proximity to the spot being deconvolved which are therefore likely to have similar cell-type composition and can be used as additional data points for inferring the cell-type proportions of a given spot. For this purpose, we modified the local multivariate GearyC (LMGC)[34] since due to the difference squared term it better captures local differences in spatially weighted transcriptomic expression (Methods). After the adapted gearyC metric for the local neighborhood, we performed permutation testing to identify the largest transcriptomic neighborhood with significant spatial autocorrelation (Methods). We use this neighborhood and a fixed bandwidth to spatially weigh neighboring spots. Across slices, negative log alignment probabilities are used to model x-y distances between spots. Along the z-axis, we utilize a Gaussian kernel to weigh each real and imputed slice based on its relative distance from the current query slice.

Like other reference-based deconvolution approaches, SpaDecoder utilizes a cell-type annotated scRNA-seq dataset. However, most methods with a few exceptions, aggregate the scRNA-seq reference cells belonging to the same cell-type into pseudo-bulk expression profiles which are used to decompose spots and to learn the cell type proportions in each spot. This is problematic since cell-state is continuous, though cell-type annotations may be discrete. SpaDecoder directly decomposes the spatial spot expression into the product of a learnable technical or batch parameter 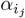, the input scRNA-seq gene by cell expression matrix *B*_*sc*_, a learnable single cell by cell-type weight matrix 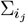 and a cell-type vector 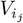 containing deconvolved cell-type proportions for the given spot *j* in slice *i* (Fig. 1f). The SpaDecoder loss function minimizes the spatial kernel weighted expression between the decomposed current spot and the expression of other neighboring spots, including the current spot, across the 3D tissue space. To avoid learning excessively large pre-softmax 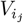 values and to enforce stability and some sparsity, we add L2-regularization.

We demonstrate the application of SpaDecoder on several downstream tasks. From the cell type proportions for each spot, we define two metrics a global (G-3DSCI) and local (L-3DSCI) 3D Cell-Type Spatial Colocalization Index to uncover cell type pair colocalization patterns for insights into cell type organization in tissue regions and tumor development. We identify tissue regions with varying proportions across slices with the Jenson-Shannon distance metric and permutation test. We perform connected component analysis to capture the spatial regions corresponding to key cell types or tumor regions. (Fig. 1g, Methods). By mapping reference scRNA-seq cells to spatial spots, we can additionally perform fine grained cell subtype annotations to identify tumor subtypes in identified regions (Fig. 1g), perform spatio-temporal location prediction, denoise the expression of measured spatial genes, and predict the expression of unmeasured genes (Fig. 1h, Methods).

### SpaDecoder improves cell-type deconvolution on three different mouse tissues

To evaluate the performance of SpaDecoder in various scenarios, we obtained single cell resolution spatial transcriptomic data from three different mouse tissues which we aggregated into square spots of fixed size determined by a parameter and comprising varying number of cells in each spot. This is preferable to synthetic (from simulated spatial data) or semi-synthetic (simulated from an scRNA-seq dataset) since the spatial locations and cell-types at each location correspond to real tissue behavior, even though single cell resolution spatial transcriptomics datasets may have fewer genes or have different resolution compared to spot based ST datasets. This gives us access to spatial cell annotations from the original publications which we use as ground truth for evaluation. To obtain a 3D stack of tissue slices, for a given sample, we simulate a stack of 10 slices, using a parameter *N*_*swap*_ to account for the dissimilarity between slices which can be the result of a large spatial distance along the z-axis or a tissue that rapidly changes in that direction (Methods). We compare SpaDecoder against top-performing deconvolution methods as identified by benchmarks - CARD [11], Tangram and the single cell reference resolution version of Tangram (Tangramsc) [6] and Cell2location[13] using three established metrics [21]: root mean square error (RMSE), Pearson correlation, and Jenson-Shannon divergence (JSD). The datasets are as follows: (a) `Moffitt2018` [35] is profiled using MERFISH on the mouse hypothalamic preoptic region comprising 132 slices (12 samples with a 10-slice simulated stack added on for each) (b) `Choi2023` [36] comprises MERFISH on the mouse retina comprising 275 slices (25 original samples) [36] and (c) `Haviv2025` [37] comprising 11 Xenium slices (1 original sample) on a mouse model of Leptomeningeal melanoma metastasis. We obtain reference scRNA-seq datasets of the corresponding tissue regions from Moffitt *et al* [35] (25,327 cells), Li *et al* [38] (20,155 cells), and Haviv *et al* [37] (9,870 cells) respectively (Supplementary Figure 6). Further information on the datasets is in Supp. Table 1.

SpaDecoder improves cell-type deconvolution on all three datasets and across all three metrics with high neighboring slice similarity or fewer cell swaps per neighborhood between successive slices (*N*_*swap*_ = 2) (Fig. 2a) as well as with lower slice similarity (*N*_*swap*_ = 5) (Fig. 2b), *N*_*swap*_ = 10 and 20 (Supplementary Figure 7). We use moscot[33] for alignment between slices (Supplementary Figure 8a). For an adjacent slice pair, as the distance between the slices increases, we expect each spot to be probabilistically aligned to more than one spot in an adjacent slice and therefore have a larger neighborhood. However, if adjacent slices are too dissimilar we desire that no neighborhood spots are selected from the adjacent slice. SpaDeocder’s neighborhood selection strategy recapitulates this, with more simulated cell swaps between slice pairs (indicative of slice dissimilarity) leading to more neighbors selected but also resulting in an increase in the spots with no neighbors from the adjacent slice (Supplementary Figure 8b).

**Fig. 2.**
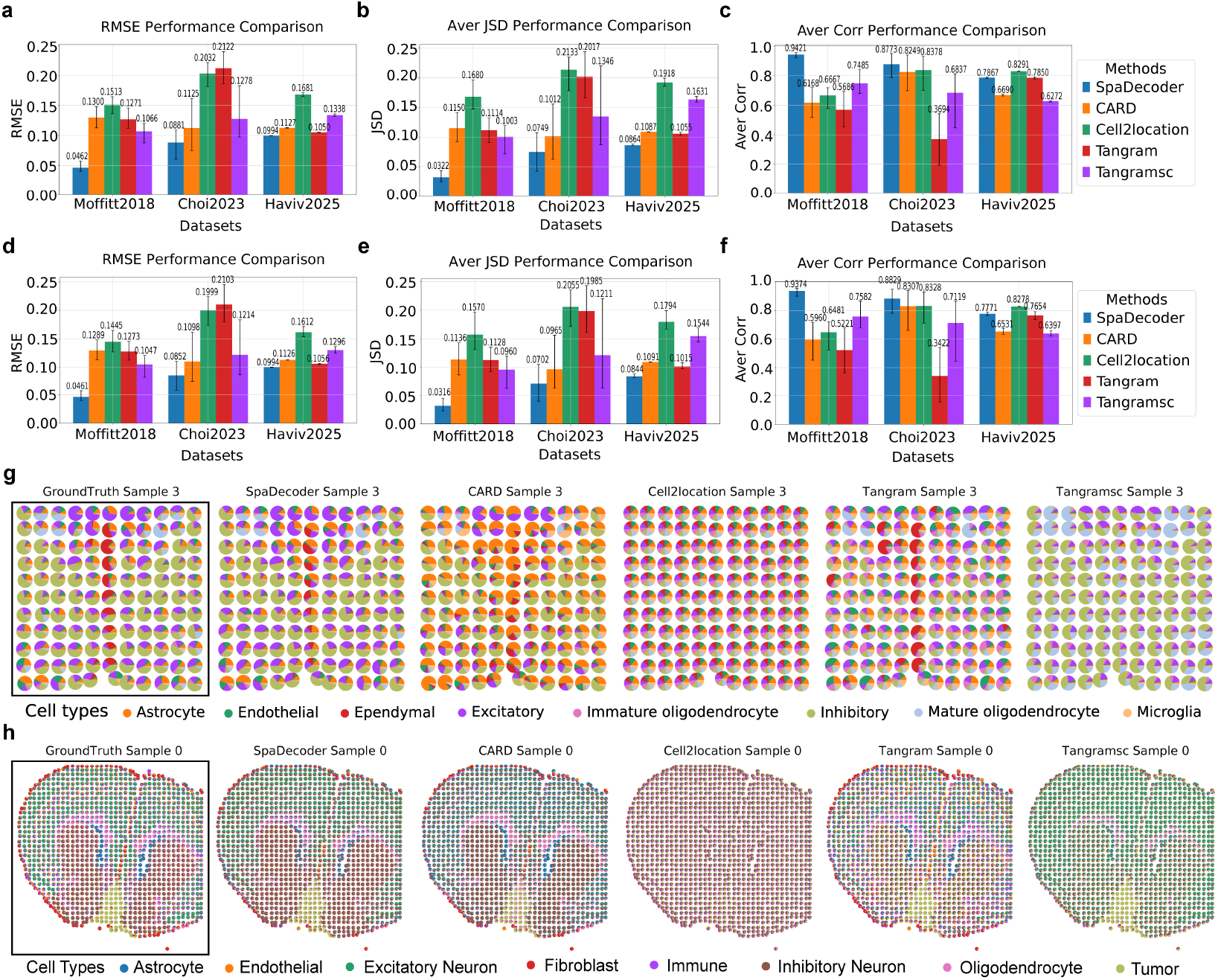
SpaDecoder qualitatively and quantitatively improves cell type deconvolution. (a-f) Barplots showing the average RMSE (a,d) JSD (b,e) Pearson correlation (c,f) of SpaDecoder, CARD, Cell2location, cluster-averaged Tangram and single cell resolution Tangram (Tangramsc) across 3 single cell spatial datasets `Moffitt2018`, `Choi2023`, and `Haviv2025` with simulated spots (spot size parameter *N*_*spot*_ = 50) and simulated slice stacks *N*_*swap*_ = 2 (a-c) and *N*_*swap*_ = 5 (d-f) (g-h) Pie plot visualizations of cell type proportions obtained from original annotations (ground truth) in `Moffitt2018` (g) and `Haviv2025` (h) alongwith deconvolved outputs from SpaDecoder, CARD, Cell2location, Tangram, and Tangramsc for representative samples (sample 3) (g) and (Sample 0) (h). Each pie corresponds to a simulated spot with colors indicating the proportion of celltypes in that spot.

Qualitative evaluation of pie plot visualizations of deconvolved cell types in each spot from the original published annotation (Ground truth), SpaDecoder, CARD, Cell2location, Tangram, and Tangramsc (Fig. 2g-h, Supplementary Figure 6) reveals that SpaDecoder most closely resembles the ground truth compared to other methods in two representative samples from `Moffitt2018` (Fig 2g, Supplementary Figure 6), a representative sample from `Haviv2025` (Fig. 2h) and two representative samples from `Choi2023` (Supplementary Figure 6). In `Moffitt2018`, compared to other methods, SpaDecoder correctly identifies the presence of the central vertical band of ependymal cells (in red) in Sample 3 (Fig. 2g) and its absence in Sample 11 (Supp. Fig. 6d) with lower false positive proportions of other cell types, an example of its effectiveness at capturing cell types at the interface between different cell type regions. Notably, ependymal cells are rare and comprise less than 0.24% of reference cells (Supp. Fig. 6a). SpaDecoder also performs qualitatively better than baselines with the retina, which has a more complex tissue tissue shape (Supp. Fig. 6e-f). Compared to other methods, pie plot visualizations qualitatively indicate that SpaDecoder distinguishes tumor (light green) from healthy tissue capturing heterogeneity in the boundary regions surrounding the tumor which is crucial for pinpointing and clinically intervening with tumor microenvironment targeted therapies (Fig. 2h).

### Ablation tests

We validate the design choices in SpaDecoder with some ablation tests. We observe that sharing information across multiple slices improves performance over single slice deconvolution (Supplementary Figure 1). Further, augmenting the query spatial stack by utilizing the alignment probabilities to linearly impute gene expression at intermediate spatial locations along the *z*-axis also improves performance (Supplementary Figure 2). Further, our adaptive neighborhood identification approach improves deconvolution metrics, confirming its utility (Supplementary Figure 3). We observe a significant improvement in performance with learning 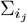 and utilizing the single cell resolution reference as opposed to pseudo-bulk reference profiles (Supplementary Figure 4). SpaDecoder also excels across varying spot size and number of cells in a spot (Supplementary Figure 8) compared to other methods, though all methods under consideration perform worse as spot size decreases which is consistent with previous findings [23]. We also find that SpaDecoder is stable within a range of parameters for regularization *λ*, 3D bandwidth, and the number of slices imputed (Supplementary Figure 9) so we keep these fixed at *λ* = 0.1, *BW* = 0.01, *N*_*slice*_ = 10 respectively for all experiments.

### SpaDecoder decodes spot composition on multiple samples of the mouse hypothalamic preoptic region

The mouse hypothalamus has been well studied due to the intricate neuronal layout, the association of specific neuronal cell-types with functional roles and the likelihood of neuronal activity being somewhat conserved between mouse and human[39]. We utilize spatial data from Moffitt *et al* [35] comprising 12 evenly spaced slices or samples along the anterior to posterior axis of the mouse hypothalamic preoptic region (1.8 by 1.8 by 0.6 mm, Bregma +0.26 to –0.34) (Fig. 3a) and 25,327 scRNA-seq cells (Fig. 3b) from a similar region to evaluate the performance of SpaDecoder. We group cells into spots (Methods) and use published annotations [35] of cell types in spots as ground truth. SpaDecoder applied to the 12 samples excels quantitatively across average RMSE, correlation, and JSD, outperforming CARD, Cell2location, Tangram and Tangramsc. Qualitatively, too, celltype patterns captured by SpaDecoder from anterior (sample 11) to posterior (sample 0) more closely resemble ground truth compared to other tools (Fig. 3c, Supplementary Figure 10). SpaDecoder recovers all reference scRNA-seq celltypes namely, astrocytes, endothelial, ependymal, excitatory and inhibitory neurons, immature and mature oligodendrocytes, and microglia with varying anteroposterior proportions (Fig. 3d).

**Fig. 3.**
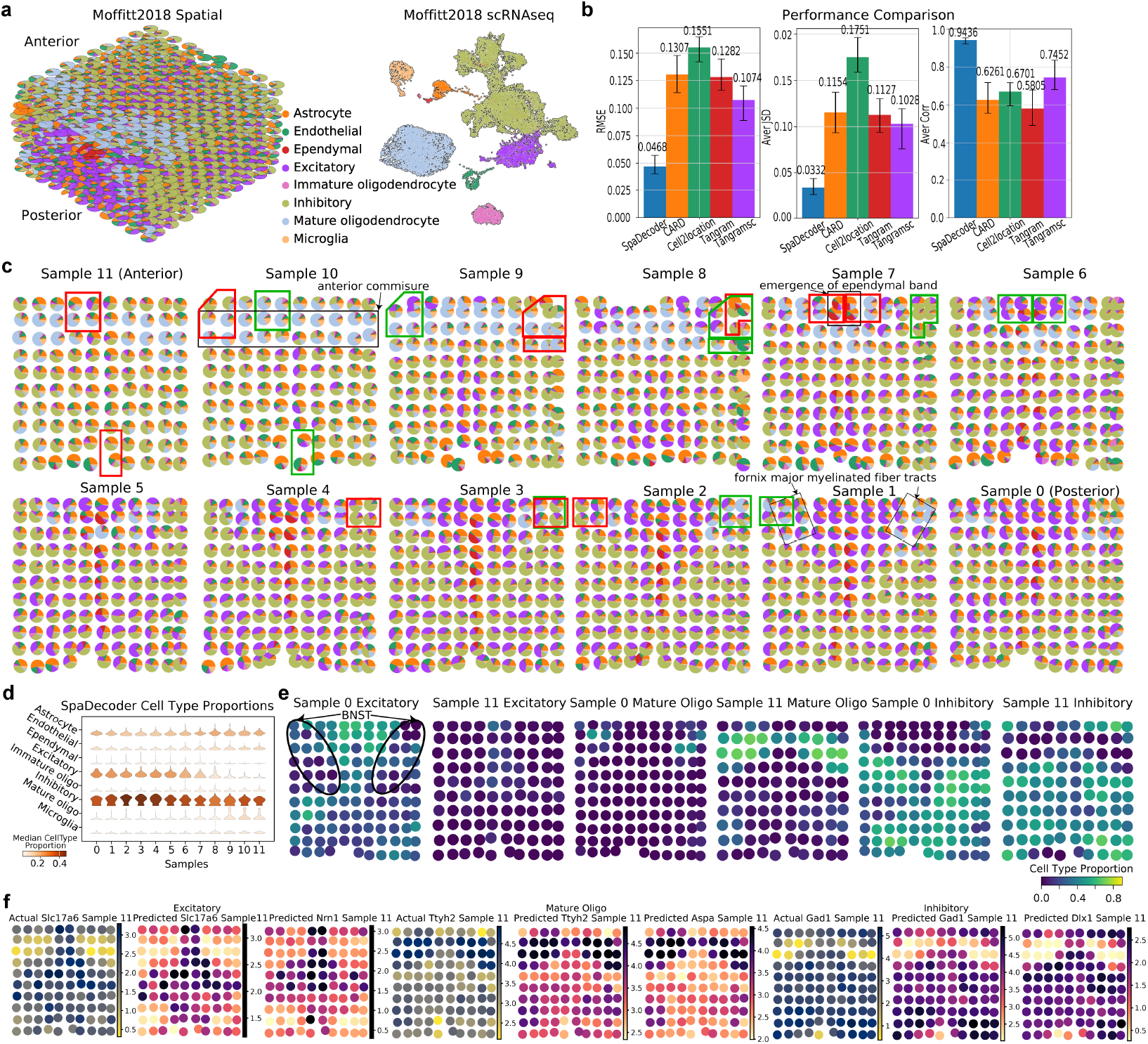
SpaDecoder captures cell type variation and expression patterns along anterior-posterior (AP) axis of the mouse hypothalamic preoptic region. (a) 3D pie plots of 12 `Moffitt2018`[35] samples from anterior (top) to posterior (bottom) with per-spot pies colored by ground-truth cell type proportions. Spots were simulated from MERFISH [35] (left) UMAP of scRNA-seq reference cells colored by Moffitt *et al* [35] annotations (right) (b) Barplots showing the average RMSE (left), JSD (middle), and Pearson correlation (right) between ground truth and outputs from SpaDecoder, CARD, Cell2location, cluster-averaged Tangram, and single-cell Tangram (Tangramsc) (c) Pie plots of SpaDecoder-estimated cell type proportions with overlaid boxes indicating regions with significant composition shifts (p-value *<* 0.1) relative to neighboring samples (posterior: red, anterior: green), computed via Jenson-Shannon distance with within-sample permutations for p-values. The samples were divided into a 6 × 6 grid and testing was performed between corresponding regions of the grid across adjacent samples. Relevant regions in accordance with Moffitt et al[35] are overlaid with black boxes and labeled. (d) Stacked violin plots showing SpaDecoder cell type proportions across samples (posterior → 0 anterior 11) (e) Spatial visualizations of posterior sample 0→ and anterior sample 11 with spots colored by SpaDecoder proportions of excitatory, mature oligodendrocytes and inhibitory celltypes. The bed nucleus of the stria terminalis (BNST) region[35] is overlaid with a black boundary and labeled. (f) Spatial visualizations of anterior sample 11 capturing log2 normalized expression patterns of known excitatory, mature oligodendrocyte and inhibitory markers measured in the experiment (leftmost in each triplet), denoised from SpaDecoder cell spot mapping (middle in triplet), and unmeasured prediction (rightmost in triplet)

In agreement with Moffitt *et al*, SpaDecoder detects the emergence of a band of mature oligodendrocytes (light blue) in the anterior commissure which are restricted to the fornix–major myelinated fiber tracts in more posterior samples [35] (Fig. 3c labeled black boxes, Fig. 3d-e). Conversely, immature oligodendrocytes (pink), astrocytes (orange), microglia (light orange), and endothelial (green) cells were scattered throughout the samples. Inhibitory (light green) and excitatory (purple) neurons exhibited distinct spatial patterns with the former being more widely dispersed [35] with higher proportions in specific regions toward the posterior (samples 2-4) while the latter are sparse in the anterior, generally more ubiquitous in the posterior but depleted in the posterior BNST (Fig. 3c-d, Fig. 3e labeled region). By automatically detecting regions changing in cell type composition with Jenson-Shannon distance and permutation testing, in agreement with anatomical findings [35], we are able to quantitatively pinpoint the start of the emergence of a band of ependymal cells (red) beginning around sample 7 and becoming more prominent toward the posterior (Fig. 3c labeled black box, Fig. 3d-e).

Leveraging SpaDecoder’s “cell to cell-type” association matrix for each spot, we obtain a reference scRNA-seq to spatial mapping matrix (Methods) and use it to denoise the expression of measured spatial genes and predict unmeasured gene expression. Visualization of key cell type specific literature-driven markers[35] across excitatory (*Slc17a6*), mature oligodendrocyte (*Ttyh2*) and inhibitory (*Gad1*) cell types (Supplementary Figure 10c) reveals high similarity between actual and denoised expression of the same gene. Correspondingly, similar patterns also arise from the prediction of unmeasured genes *Nrn1* [40], *Aspa*[41], and *Dlx1* [42] (Fig. 3f) illustrating the utility of SpaDecoder at single cell resolution spatial mapping for spatial gene expression denoising and prediction.

### SpaDecoder provides fine-grained molecular insights in human breast cancer tissue

Breast cancer can be classified into *in situ* (ductal and lobular) and invasive [43] based on localization. Ductal carcinoma in situ (DCIS) is non-invasive or pre-invasive stage 0 breast cancer and often progresses to invasive ductal carcinoma (IDC) [44]. However, the initiation of DCIS and progression to invasive is poorly understood [44]. Therefore, automatic fine grained localized spatial understanding of breast cancer tissue regions, associated cell types and underlying molecular characteristics could prove extremely beneficial for diagnosis and treatment.

Here, we apply SpaDecoder to two 5*µm* breast cancer tissue samples or slices along with a reference scRNAseq dataset comprising 26, 031 cells from Janesick *et al* [45] (Fig. 4a-b). The spatial samples were profiled at single cell resolution with 10x Xenium and 3201 and 2199 spots were obtained respectively using our spot generation procedure (Methods). Ground truth cell type proportions for the spots were obtained from given annotations in Janesick *et al* [45]. SpaDecoder identifies B, dendritic (DCs), Endothelial, Macrophages, Mast, Myoepithelial (ME), Perivascular-like, Stromal, T, and Tumor cells in both samples with patterns resembling the corresponding sample ground truth according to the original published annotations (Fig. 4a). Qualitatively, our identified cell-type patterns more closely resemble ground truth compared to CARD, Cell2location, Tangram and Tangramsc (Fig. 4a, Supplementary Figure 11a-d) and quantitative evaluation across Average RMSE, Pearson Correlation, and JSD confirms SpaDecoder’s superior performance over baseline methods (Fig. 4c). Visualizing the proportions of the key cell types (tumor, stromal, T, and ME) revealed spatial patterns in the cell-type distribution (Fig. 4d). In particular, we observe a large mostly connected region of stromal cells, several groups of tumor cells on left and right sides with different spatial shapes, and some less pronounced grouping of T and ME cells.

**Fig. 4.**
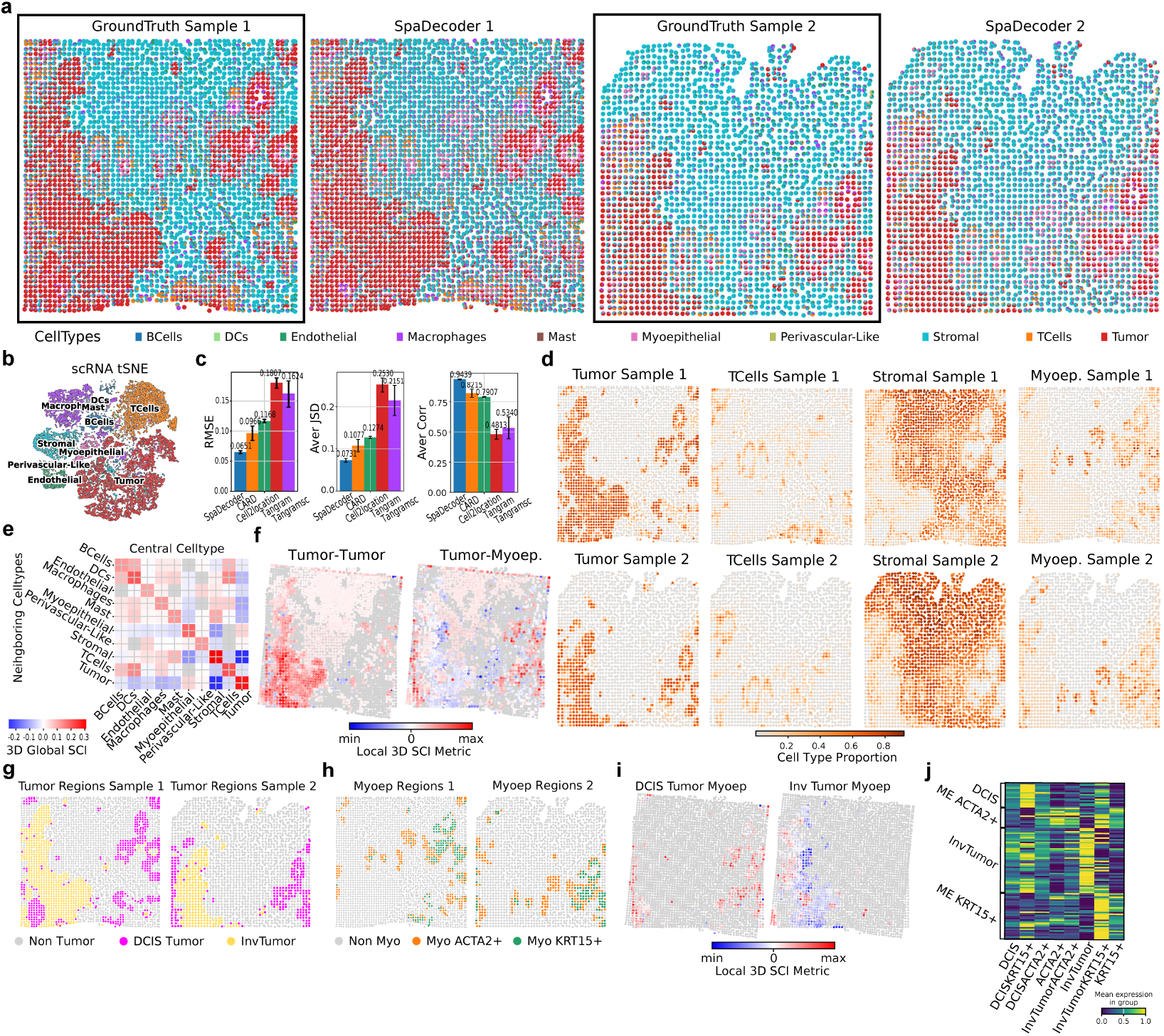
SpaDecoder decodes cell types and uncovers molecular insights in human breast cancer tissue. (a) Pie plots of cell type proportions from ground-truth annotations (Janesick et al[45], black box) and SpaDecoder on two Xenium samples with neighborhood-aggregated spots (b) tSNE embedding of scRNA-seq reference cells (Janesick et al [45]), colored by cell type (c) Barplots showing average RMSE, JSD, and Pearson correlation between ground truth and deconvolved proportions from SpaDecoder, CARD, Cell2location, Tangram, and Tangramsc (d) Spatial plots of sample 1 and 2 with spots colored by SpaDecoder proportions of key celltypes (Tumor, T, Stromal, and Myoepithelial (ME)) (e) Heatmap of global averaged 3D local neighborhood colocalization between cell type pairs (central type: X-axis; neighbor type: Y-axis). Negative values indicate neighboring type depletion, positive values enrichment; nonsignificant pairs (p¿0.05) in gray. Significance is obtained via permutation testing (Methods) (f) Aligned slices showing 3D local colocalization for Tumor-Tumor and Tumor-Myoepithelial (ME) cell type pairs. The first in the pair indicates the central celltype while the second is the neighborhood celltype. Negative values indicate neighboring type depletion, positive values enrichment; nonsignificant (p-value > 0.05) in gray (g) Spatial slices colored by tumor subtypes identified from tumor regions obtained from connected component analysis of SpaDecoder tumor celltype proportions (h) Spatial slices colored by ME subtypes identified from ME regions obtained from connected component analysis of SpaDecoder ME celltype proportions (i) Aligned samples showing 3D local colocalization for DCIS–ME, and invasive tumor–ME; nonsignificant spots (p-value > 0.05) in gray. (j) Heatmap of scaled mean expression of data-driven, cell type–enriched scRNA-seq markers (Y-axis) across tumor and ME subtype spatial regions visualized in Supplementary Fig. 11f.

We then investigated if any of the cell types had strong colocalization. Utilizing the SpaDecoder cell-type proportions, we computed our Global 3D Cell-Type Spatial Colocalization Index (G-3DSCI) (Methods) (Fig. 4e). We observe that tumor cells are anti-colocalized with the broadly grouped stromal cells but positively colocalized with other tumor and ME cells (Fig. 4d-e). This means that compared to a random 3D neighborhood, on average, tumor cells are more likely to occur in the immediate vicinity of other tumor and ME cells and less likely to co-occur with healthy stromal cells. The colocalization of ME and tumor cells is interesting since the former have been previously shown to play a critical role of tumor progression[46]. We therefore sought to localize spots where tumor cells were surrounded by neighboring cells significantly (*pval <* 0.01) enriched or depleted in ME cells. To this end, we used our L-3DSCI metric (Methods) which tested the colocalization in the neighborhood surrounding each spot compared to a randomly chosen background. The results revealed individual spots with neighborhoods of colocalization of tumor cells with each other, and tumor cells with ME cells (Fig. 4f). Since tumor cells often group together in specific patterns and are surrounded by other cell types, we performed cell type region identification (CTRI) to spatially localize contiguous tissue regions containing cells of a particular type with connected cell component analysis (Methods) and identified T, tumor and ME regions (Supplementary Fig. 11e). Leveraging fine grained annotations from Janesick *et al* [45] and our single cell to spatial mapping, we further classify tumor and ME regions into their respective cell subtypes using “cell subtype identification in regions” (Methods) and identify DCIS tumor, Invasive tumor, ACTA2+ ME, and KRT15+ ME cells (Fig. 4g-h, Supplementary Fig 11f). We observe a distinct pattern of invasive cells to the left in both samples while DCIS cells occur on the left and right but have a more pronounced oval shape toward the right, consistent with Janesick *et al* [45]. Further, unlike on the left where ME cells are fewer and less structured, ME cells on the right form roughly concentric ME KRT15+ (inner) and ME ACTA2+ (outer) patterns. We also note the existence of ACTA2+ regions without the presence of KRT15+ regions, a pattern also observed in Janesick *et al*. It also appears that DCIS colocalizes with ME, which is confirmed by L-3DSCI (Methods) analysis showing significant positive colocalization between DCIS and ME with almost no negative colocalization on both left and right sides of the samples (Fig. 4i). Additionally, we observe few positively and largely negatively colocalized Invasive tumor and ME cell spot neighborhoods (Fig. 4i). This indicates a significant presence of ME cells in DCIS neighborhoods and an absence in most invasive neighborhoods, consistent with the hypothesis that ME cells act as a barrier and loss of ME cells and their associated basement membrane is a hallmark of the transition from DCIS to IDC[46]. We confirmed our DCIS, invasive, ACTA2+ ME, KRT15+ ME cell subtype annotations by identifying differentially expressed genes in the corresponding categories from the reference scRNA-seq dataset (adjusted p-val *<* 0.05, log2FC *>* 1.0) and visualizing their expression on the spots with annotated subtypes or pairs of subtypes with good consistency (Fig. 4j).

In conclusion, SpaDecoder qualitatively and quantitively proved successful in identifying cell types. Utilizing SpaDecoder outputs, we performed downstream analyses and identified cell subtypes, uncovered celltype occurence and colocalization patterns globally as well as at spot resolution. Further, we identified putatively interesting regions of tumor and ME cooccurence, annotated their subtypes and provided evidence supporting the hypothesis that ME cells act as a barrier to the transition from DCIS to invasive tumor[46].

### SpaDecoder reveals spatio-temporal cell type organization during human fetal thymic development

SpaDecoder has demonstrated excellent performance across various tissues and technologies by sharing information across 3D spatial tissue slices. We next inquire if SpaDecoder can perform well in the spatio-temporal scenario, such as during organ development when 2D tissue slices are profiled across time. We sought a recently published spatial atlas capturing human fetal and pediatric thymic development [47] and selected the longest continuous range of spatially profiled time points comprising 4 10X Visium samples spanning 16 to 19 post conception weeks (pcw) sampled weekly and scRNA-seq data at 15pcw, 17pcw, and 18pcw.

The thymus is a specialized organ of the immune system responsible for T Cell maturation and education[48]. It has two lobes joined by connective tissue and surrounded by a connective tissue capsule with septae or partitions that extend inward from the capsule[47]. Internally, it has two distinct compartments, the outer dense cortex and inner medulla, along with secondary structures[49]. Tcell maturation which begins as early as 8 post conception weeks (pcw) in humans is a highly organized and regulated process[50]. Developing Tcells are broadly categorized based on their expression of CD4 and CD8 into CD4-CD8-Double negative (DN), CD4+CD8+ double positive (DP), or single positive (SP) (either CD4+CD8- or CD4-CD8+)[51]. Early Tcell Progenitors (ETPs) enter the thymic medulla around the corticomedullary junction (CMJ) and migrate to the cortex where they may undergo maturation to DN and SP through interaction with cortical thymic epithelial cells (cTECs), differentiate into CD4+ and CD8+ T cell lineages via positive selection and migrate to the medulla [52].

After excluding ambiguous annotations, we were left with 34,798 cells visualized on the published UMAP embedding (Fig. 5a) colored by annotated cell type (top) and developmental day (bottom) [47]. We selected an scRNA-seq annotation resolution that would capture the heterogeneity in major cell types but was not overly fine-grained and indiscernible. In particular we annotated Schwann cells, red blood cells (RBCs), myeloid, cTECs, mTECs, mcTECs (putative bipotent TEC progenitor cells), mimetics, DN, DP, and SP cells from the original published annotations [47] (Fig. 5a, top). Despite the variability in spatial tissue shape across the samples, SpaDecoder identified all the cell types (Fig. 5b) qualitatively, based on visual similarity to known thymic structure described above, outperforming Cell2location and Tangramsc (Supplementary Figure 12). Visualization of deconvolution proportions on the spatial embeddings at each of the time points revealed key regions. At each time point, consistent with thymic structure, we identify stromal cells in the capsule and along the septae (Fig. 5c). DN cells are in lower proportions as expected but concentrated around the periphery consistent with their presence in the cortex. DP cells are much more prominent than DN but also largely in the cortex, unlike SP cells which are highly concentrated in the interior medulla. As expected, the thymus is overrun with developing T-lymphocytes and TECs are much less abundant (Fig. 5c). Therefore, to further delineate mTEC and cTEC regions, we performed cell type region identification (Methods) and found that while cTECs are present in the outer cortex, the mTECs are in higher proportion with more clearly demarcated localization patterns in the inner medulla (Fig. 5c-d). Interestingly, the cortex is subdivided into more regions than the medulla which could be due to the septae that extend inward from the capsule partially fragmenting it (Fig. 5c-d). Despite the fact that the two thymic lobes are clearly demarcated at 16 and 19pcw while they are less clear in the 17-18pcw samples likely due to technical differences when processing, SpaDecoder is able to capture the cell-types effectively (Fig. 5b-d).

**Fig. 5.**
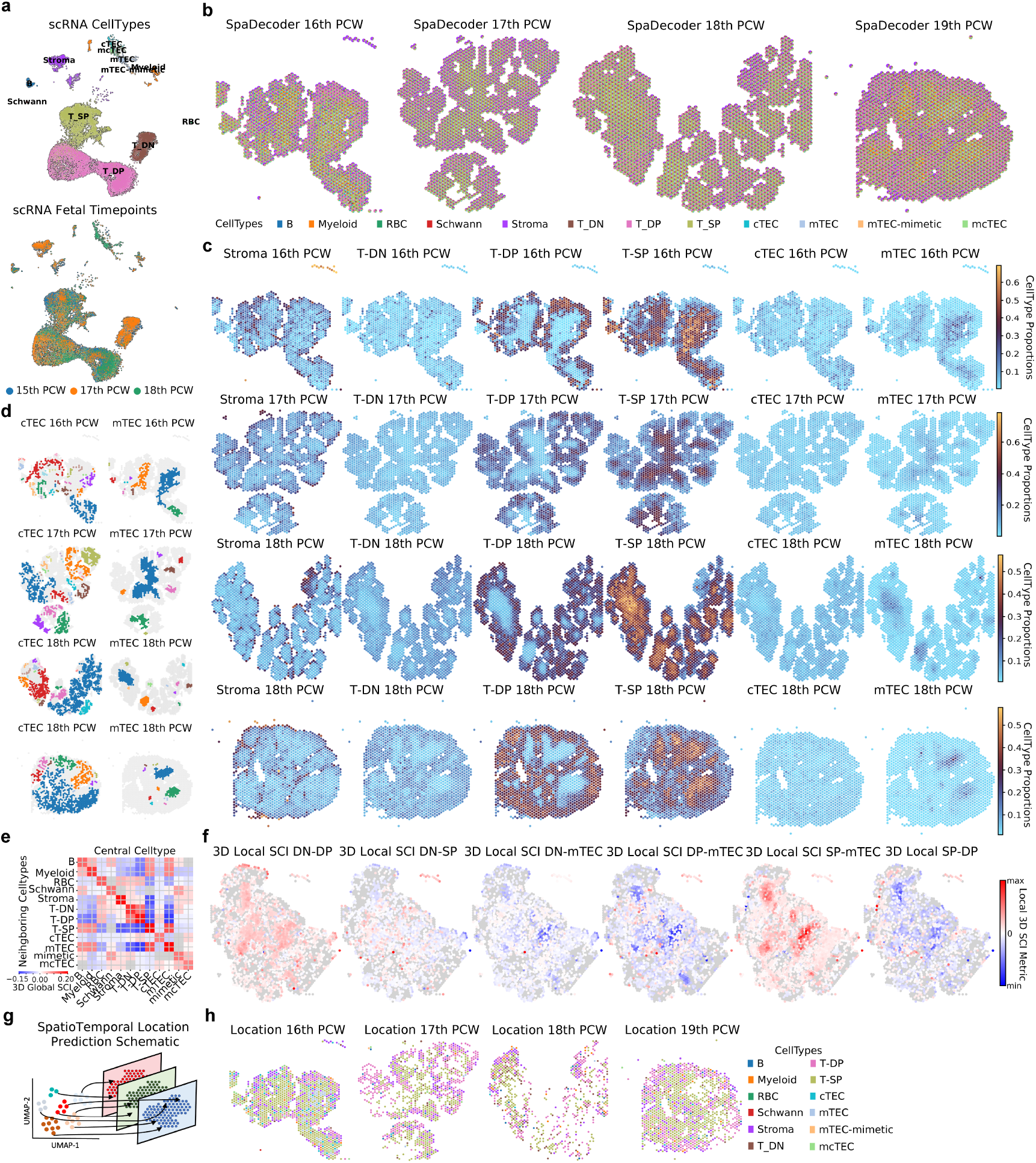
SpaDecoder decodes spatiotemporal tissue organization during human fetal thymic development. (a) UMAP of scRNA-seq reference cells (Yayon et al [47]) colored by cell type (top) and fetal timepoint (bottom) (b) Pie plots of SpaDecoder-inferred cell type proportions on Visium slices or samples from 16-19PCW of human fetal thymic development (c) Spatial plots showing SpaDecoder proportions of key cell types (Stroma, DN-T, DP-T, SP-T, cTEC, mTEC). (d) cTEC and mTEC regions identified at each timepoint via connected component analysis (e) Heatmap of averaged 3D local neighborhood colocalization between cell type pairs (central type: X-axis; neighbor type: Y-axis); negative values indicate depletion, positive enrichment; nonsignificant pairs (p> 0.05) in gray. (f) Aligned samples showing 3D local colocalization for DN-DP, DN-SP, DN-mTEC, DP-mTEC, SP-mTEC, and SP-DP; nonsignificant spots (*p* > 0.05) in gray (g) Schematic of SpaDecoder-inferred cell-to-spot mapping matrix predicting single-cell spatiotemporal coordinates (h) Predicted spatial coordinates of reference scRNA-seq cells across timepoints colored by cell type.

Consistent with the described structure of the thymus, global and local cell type colocalization revealed positive spatial colocalization between DN and DP in the cortex, and between SP and mTECs which are both medullary (Fig. 5e-f). We also observed negative spatial colocalization between DN (cortical) and SP (medullary), DN and mTEC, between DP and mTECs, and between SP (medullary) and DP (cortical) (Fig. 5e-f), consistent with thymic structure. Finally, we recapitulate the spatio-temporal cell layout by mapping each scRNA-seq cell to the spot across time and space with the maximum mapping probability (Fig. 5g) and observe that the recovered spatial patterns closely resemble the original spatial layout at each time point (Fig. 5h).

## Discussion

In this work we present SpaDecoder, a spatial spot deconvolution method leveraging individual single cell reference profiles, as opposed to cell-type aggregated, and 3D spatial tissue structure with the help of an adaptive 3D weighted spatial Gaussian kernel which enables information sharing across transcriptionally similar spatially proximal spots. We predict cell-type proportions by optimizating a matrix factorization based loss function. In order to adapt to both homogeneous as well as heterogeneous tissue environments, we develop a permutation based localized weighted spatial autocorrelation metric which when applied to each spot efficiently selects the neighborhood within which to pool transcriptomic and spatial information to improve deconvolution predictions. We probabilistically align adjacent tissue slices in 3D and infer intermediate slice expression to augment our 3D spatial tissue slice stack. Utilizing a learnable parameter, we model the batch effect between the reference scRNA-seq and the spatial dataset. We extensively evaluate SpaDecoder in several scenarios. To overcome the lack of ground truth, we first utilize simulated spots from single cell spatial transcriptomic datasets and simulate tissue slice stacks by perturbing a single slice. We use three datasets encompassing various tissue types: MERFISH on the mouse hypothalamic preoptic region from Moffitt *et al* [35], MERFISH on the mouse retina from Choi *et al* [36] and Xenium on a mouse model of Leptomeningeal melanoma metastasis from Haviv *et al* [37]. We present quantitative results with established evaluation metrics (root mean square error, Jenson-Shannon Divergence, Pearson correlation) utilizing the original published annotations from spatial single cells as the ground truth. SpaDecoder excels with varying spot sizes and slice stack simulation conditions. While we chose moscot [33] for slice alignment due to good performance, any alignment method may be used provided it can estimate probabilistic mapping between slices and allow imputation of intermediate slices between every pair. SpaDecoder is stable within a range of hyperparameter choices so dataset specific tuning is unnecessary. Detailed ablation tests reveal improvement with the use of single cell resolution reference scRNA-seq data, multiple spatial slice deconvolution, intermediate slice imputation for data augmentation, and our adaptive neighborhood selection strategy.

SpaDecoder qualitatively and quantitatively improved deconvolution on 12 anterior-posterior (A-P) MER-FISH samples of the mouse hypothalamic preoptic region from Moffitt *et al* [35] aligned using moscot[33]. We identify cell types with characteristic spatial organization patterns, and uncover regions that gradually change composition along the AP axis in 3D stack. We map scRNA-seq cells to spatial spots enabling denoising of measured spatial genes, and the prediction of unmeasured markers associated with key celltypes. On 2 breast cancer samples profiled by Janesick *et al* [45], SpaDecoder identifies distinct tumor, stromal, T and myoepithelial (ME) cell patterns, and spatially colocalized cell types across 3D. We categorize tumor regions into ductal carcinoma *in situ* (DCIS) and invasive, and ME regions into ACTA2+ and KRT15+. Our analyses supports the hypothesis that ME cells act as a barrier for progression from DCIS to invasive tumor[46].

SpaDecoder additionally excels on a 3D spatio-temporal human fetal spatial thymic atlas profiled with 10X Visium capturing 16pcw-19pcw sampled weekly with a reference scRNA-seq atlas across 15pcw, 17pcw, and 18pcw. Despite the technical variations in tissue shape, SpaDecoder performed qualtitatively well, capturing celltypes, global and local colocalization patterns, and regions according to their known occurence in the cortical and medullary thymic regions. Importantly, SpaDecoder spatio-temporally mapped scRNA-seq cells to spatial spots showing broad application potential to spatially profiled developmental and disease time courses.

## Methods

### SpaDecoder

Given input data comprising 3D spatial transcriptomic data capturing the transcriptomic profiles of measured genes alongwith their 3D spatial coordinates (or 2D coordinates and relative spatial or temporal positions along the z-axis), spatial deconvolution delineates the proportion of different celltypes at each location. The complexity lies in the fact that each coordinate location contains a collection of varying number of single cells termed “spots”. SpaDecoder, like several other tools, [6][13][11], leverages a separately profiled reference scRNA-seq dataset of similar tissue type with predefined celltype annotations. To preserve the cell-cell heterogeneity, unlike other approaches, SpaDecoder decomposes the spatial spot gene expression into the product of reference cell expression profiles, a learnable assignment of reference cells to predefined celltypes and a learnable cell type proportion vector. We formulate the objective as a matrix factorization problem and solve it separately for each spot with a highly parallelized implementation that speeds up runtime. Additionally, SpaDecoder utilizes cell type expression profiles from 3D neighboring spots (within and across slices) which share similar expression profiles and therefore likely have similar cell type composition. Since the homoegeneity of spot neighborhoods depends on the microenvironment, we additionally define a metric to optimally select the relevant 3D neighboring spots. When utilizing a 3D stack of 2D tissue slices, the individual slices are misaligned due to technical artifacts or varying tissue shape. We align slices with an optimal transport based strategy [33] and further infer intermediate slices to bridge the gap between adjacent slices and augment the dataset. We apply an additional spatially decaying kernel along the z-axis to account for increasing dissimilarity between slices that are spatially or temporally far apart. We describe these innovations and our model here with further details in the Supplementary file. The workflow of SpaDecoder follows Figure 1.

#### Objective Function

We denote the gene expression count matrix of the spatial transcriptomic dataset as **X**, where the expression of spot *j* (*j* = 1, 2, · · ·, *N*_*spa*_) is the column vector **x**_*j*_, and denote the count matrix within scRNA-seq dataset as **B**_**sc**_, where the expression of cell *m* (*m* = 1, 2, · · ·, *M*) is the column vector **b**_*m*_. Let *G* be the number of genes common to both the scRNA-seq reference and the spatial dataset. SpaDecoder decomposes the expression of spot *j* following the Mean Square Error (MSE) objective:

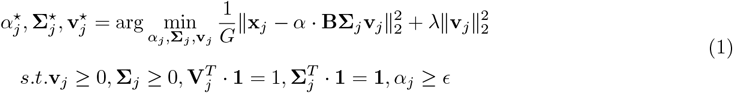

For each spot *j*, we learn a cell-by-cell-type label association matrix **Σ**_*j*_, scalar *α*_*j*_, and cell-type deconvolution scores **v**_*j*_. **Σ**_*j*_ and **v**_*j*_ are further constrained to be unit simplex using softmax which better resembles the compositional proportions. 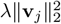 is the regularization term of **v**_*j*_. By constraining the columns of **Σ**_*j*_ to a unit simplex we ensure all, even rare celltypes, are equally weighted in the scRNA-seq reference. This is particularly useful if the reference and spatial datasets differ in their celltype composition. In such cases, weighting celltypes with reference based priors would be disadvantageous. Further, the learned association matrix can assign varying total (across celltypes) weights to each cell which encourages higher weights for more representative cells and lower weights for noisy cells. When deconvolving spot *l* in slice *k*, to further utilize 3D spatial information, SpaDecoder considers not only the expression value 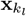 of spot *l*, but also the expression values 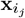, for all spots *j* in slice *i* in the 3D the spatio-transcriptomic neighborhood 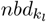 of spot *i*_*j*_:

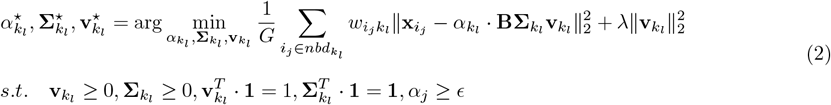

The final objective function is the weighted sum considering all *l*’s transcriptionally similar spatially proximal spots *j*, and the weight 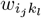 represents the 3D similarity between spot *j* and spot *l*. 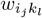 captures transcriptomic similarity as well as the relative 3D spatial locations of spots. Since each spot is optimized separately, we speed up runtime by computing in parallel on a GPU the deconvolution scores for all spots (or optionally a fixed user provided batch size) in the 3D tissue. Further, each spot can have variable number of neighbors depending on whether it is located in a homogenous or heterogenous region. The final output of SpaDecoder includes, for each spot *k*_*l*_, the cell-type deconvolution scores 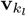, label association matrix 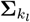 and scalar 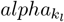. 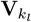 can be used directly as the cell-type decomposition scores similar to the traditional spatial deconvolution scores, where each element shows the proportion of the corresponding cell type that composes the spot *k*_*l*_ (cell-type-level decomposition). In addition, the matrix product of 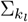 and 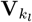 shows the proportion of each single-cell that composes the spot *j* (cell-level decomposition) which we use for downstream tasks (See Applications). Detailed notation is included in Supplementary Note 1.

#### SpaDecoder Preprocessing

##### Data Preprocessing

We apply standard preprocessing corresponding to best practices for handling scRNA-seq or spatial datasets [53]. Raw scRNA-seq count matrices were obtained from the respective datasets (described in Data section below) and standard pre-processing was performed using scanpy[31]. Genes present in less than 10 cells were excluded and the subset of overlapping genes between scRNA-seq and spatial data was selected. Cells with minimum total counts *<* 50 were excluded. In case there were over 5000 resulting genes in the set, we performed additional gene feature selection. For scRNA-seq data, we normalize (sc.pp.normalize_total) and log pseudo count transform (sc.pp.log1p) the count matrix, and select the differentially upregulated genes in each defined scRNA-seq cluster (adjusted p-value *<* 0.05, log-foldchange *>* 1.0). We concatenate all spatial slices and select the top 5000 slice-aware highly variable genes (sc.pp.highly_variable_genes) to avoid the selection of slice specific genes. The overlap between spatial highly variable genes and scRNA-seq differentially upregulated genes comprises the resulting feature set. Any feature selection approach can be used. Prioritizing genes that are meaningfully (variably) expressed in the spatial data as well as that are cell-type enriched in the scRNA-seq data will likely maximize performance.

scRNA-seq and spatial data is normalized using sc.pp.normalize_total with target sum = 1000 to correct for varying sequencing depth. Since different spatial datasets might have different coordinates and spatial distances between spots, we scale X-Y spatial coordinates to [0, 1].

##### Slice Alignment

While the user may choose to run our tool in single slice mode, SpaDecoder is primarily designed for information sharing across stacks of 2D slices. We leverage an optimal transport strategy, moscot[33], but any alignment method which infers a pairwise distance or probability between spots in adjacent slices may be used (Fig. 1e). Moscot[33] improves on PASTE[5] and outputs a coupling matrix *P* between each pair of slices where *P*_*ij*_ denotes the amount of probability mass transported from spot *i* in the left dataset to spot *j* in the right dataset. We use the balanced entropic regularized Fused-Gromov Wasserstein Optimal Transport (FGW-OT) formulation to minimizes the transcriptomic distance between mapped spots across the two slices as well maximize the correspondence between spatial arrangement of spots for each slice with its immediate neighbors. For two slices *X* and *Y*, let *a* denote the set of spots in *X* and *b* the corresponding set in *Y*, then the general form of an FGW-OT problem is

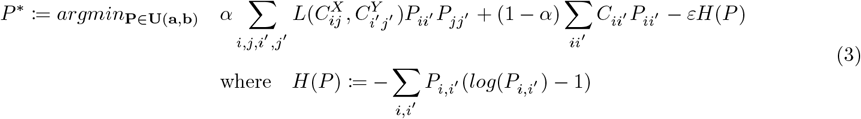

for spots *i, j* ∈ *X* and *i′*, *j′* ∈ *Y* where the first GW term quantifies the spatial correspondence between the slices, the second W term represents transcriptomic matching with *α* controlling the weight between them and *H*(*P*) is the entropy regularization term with strength *ε*. To obtain the matrix *C*, the normalized preprocessed counts from both slices are projected onto a joint PCA space (with 30 components by default) and the squared Euclidean distance is computed between pairs of spots from the two slices. The matrices *C*^*X*^ and *C*^*Y*^ are obtained from the squared Euclidean distance on normalized spatial coordinates in slice *X* and *Y* respectively. 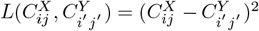. In SpaDecoder, we set *α* = 0.5 for all the experiments. The full rank solver from Peyré et al[54] as implemented in moscot[33] was used.

Interestingly we did not observe any improvement in deconvolution performance with joint iterative alignment and deconvolution, that is, by performing single slice deconvolution with SpaDecoder, incorporating the celltype proportions in the FGW-OT objective with an updated linear cost 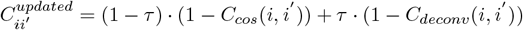 and using the resulting alignment for multi-slice deconvolution. Here *C*_*cos*_ is the cosine distance between the PCA embeddings of spot *i* and *i′*, *C*_*deconv*_ is the Pearson correlation between the deconvolution scores of *i* and *i′*, and *τ* is a weighting parameter.

##### Intermediate Slice Imputation

To add data points to our SpaDecoder deconvolution objective as well as leverage assumed variation in cell type composition along the *z* axis, we perform intermediate slice imputation (Fig. 1e). For each adjacent slice pair in the input dataset, we simulate *N*_*imp*_ slices. Let *P*_*pq*_^*ij*^ be the alignment probability between the *p*th spot in slice *i* and *q*th spot in slice *j* obtained from slice alignment. Let *t* = 1, 2, · · ·, *N*_*imp*_ − 1 be the slices to impute. **x**_*p*_^*i*^, **x**_*q*_^*j*^ is the *G* × 1 expression vector of the *p*th spot in slice *i* and the *q*th spot in slice *j* respectively. *N*_*i*_ is the number of spots in slice *i*. 𝒩 (*µ*, Σ) represents Gaussian noise with mean *µ* and standard deviation Σ. Then we perform linear interpolation to obtain expression vector **y**_**q***t*_^*j*^ for spot *q* in intermediate slice *t* which is of the same shape as slice *j*. We set *N*_*imp*_ = 20 by default but observe stable performance between 10 and 20 (Supplementary Figure 9).

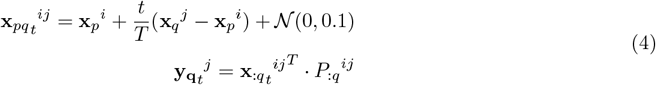

##### Adaptive Neighborhood Selection and Kernel Estimation

Let 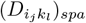 be the spatial distance between the *j*th spot in the *i*th slice and the *l*th spot in the *k*th slice. For two spots in the same slice, *i* = *k*, we define 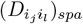 as the squared Euclidean distance. Otherwise, 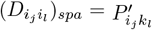 is derived from the alignment probability matrix *P* ^∗^(3). In this case, 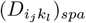 may not be symmetric since the number of spots in the two slices may be different. Let slice *i, k* have *N*_*i*_, *N*_*k*_ spots with *k* being the current reference slice and *i* the neighboring slice. Then, we scale the transport matrix 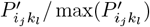, clip low transport probabilities at 0.00001, convert it into a negative log distance and rescale by the maximum value to get 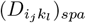. Once 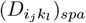 is obtained, we perform filtering 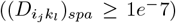, normalization 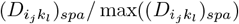 and then define an X-Y Gaussian kernel to model the weighted spatial distances between any two spots:

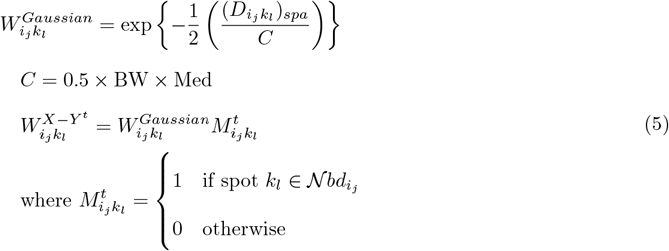

*C* is a constant determined by the hyperparameter bandwidth BW and Med is the median 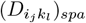 in the slice. We define *t*_*max*_ binary masks **M**^*t*^ for each spot, *t* = 1 · · · *t*_*max*_ by selecting the top *t* transcriptomic neighbors as measured by squared Euclidean transcriptomic distance 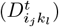 and setting low distance *<* 1*e*^−10^ to 0. If a spot lies in the neighborhood of another spot the mask is 1, else 0. For our purposes here, a spot is a neighbor of itself. We choose *t*_*max*_ = 30.

Spatially proximal spots may not always have similar transcriptomic profiles, particularly in complex heterogeneous tissue structures or at the boundary between cellular structures[29]. This is also tissue, region and even spot dependent since we would benefit from using more neighbors in more homogeneous tissue environments compared to fewer neighbors in heterogeneous regions. Hence, we perform permutation testing by defining an adapted weighted GearyC metric[55] to select the spatial neighborhood with high transcriptomic similarity for each spot *k*_*l*_ and mask *M* ^*t*^:

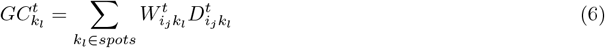

Lower *GC* values indicate lower weighted transcriptomic distances and therefore are a good choice for neigh-borhood. We perform permutation testing for each spot, by permuting all other spots, and computing *GC N* = 500 times to obtain a p-value and restrict to significant neighborhoods (p-value *<* 0.05). To maximize the number of data points available, we select the kernel from the significant neighborhood with the largest number of neighbors as the X-Y kernel 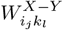. If no neighborhood has significant *GC*, the spot likely lies in a highly heterogeneous region, so we define the neighborhood to be the spot itself. This operation is parallelized across all spots, slices, and iterations, with optional selection of batch size, thereby optimizing runtime. For all augmented slices between a current *k* and neighboring slice *i*, the 2D kernel weights are **W**_**ij**_ since the X-Y spatial locations do not change with augmentation in our approach, with variation captured along the z-axis.

We further utilize a Gaussian kernel to account for the spatio-temporal variation along the z-axis. We index the real and augmented slices in the stack and For neighboring slice with index *i* and selected slice with index *k*:

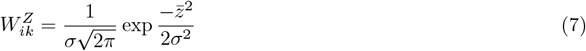

where 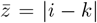 is the slice gap and *σ* = *BW*_3*D*_*/*4 where *BW*_3*D*_ is the 3D bandwidth hyperparameter. We set *BW*_3*D*_ = 8, though results are stable in 8 to 16 (Supplementary Figure 9). The resulting kernel in 2 is the product of the X-Y and Z kernels:

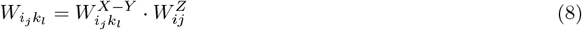

### Applications of SpaDecoder

We define several metrics from SpaDecoder outputs and use them for biological insights (Fig. 1g).

#### Global 3D Cell-Type Spatial Colocalization Index (3DC-SCI)

Utilizing the SpaDecoder cell-type proportions, we developed a 3D spatial cell colocalization metric motivated by the 2D global gene spatial cross-correlation index (SCI) in Meringue [56] (Fig. 1g). For every slice, we stack the slice before and after (except for the first and last slice which is a stack of 2), align them, and project the X-Y coordinates onto the center slice with moscot[33]. For the z-coordinate, use the the position in the tissue or an arbitrary index which preserves the ordering. We obtain the 3D spatial neighbors with Delaunay triangulation[57] and ensure that a spot is a neighbor to itself since a spot may comprise *>* 1 celltype. Then we compute 3DC-SCI for cell-types *x* and *y* as:

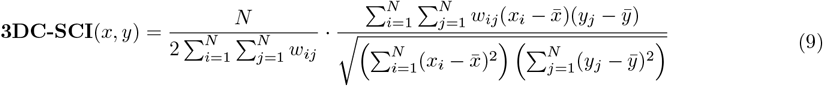

where *N* is the number of spots, *W*_*ij*_ is the binarized spatial connectivity matrix, *x*_*i*_ and *y*_*j*_ are the cell-type proportions of cell-type *x* and *y* at spot *i* and *j* and 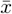 represents the mean cell-type proportion across the stack. Next, we recompute the metric after permuting the spots *P* = 1000 times to obtain a p-value.

#### Local 3D Cell-Type Spatial Colocalization Index (L3DC-SCI)

Similar to 3DC-SCI we define L3DC-SCI as the localized cross correlation between the cell-type proportion in the center spot and its neighbors as follows:

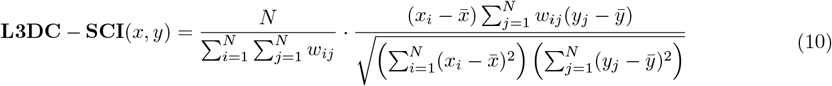

As before, we recompute the metric after permuting the spots *P* = 1000 times to obtain a p-value.

#### Cell-Type region identification (CTRI)

We group together contiguous spots containing cells of the same type into regions (Fig 1g). To accomplish this, we first classified spots based on whether or not they contained a particular cell type using the proportions from SpaDecoder, and our neighborhood smoothed binarization scheme. For each celltype *i* and spot *x* we smooth the celltype proportion across its 3D neighborhood of size *K*, filter out low proportion spots by preserving *x*_*i*_ *> T*_1_. *T*_1_ can either be explicitly defined or can be selected automatically using Otsu’s method for image thresholding [58]. The latter finds the threshold that optimally separates the two classes (presence or absence of the celltype) by maximizing the between class variance and minimizing the within class variance of the proportion of the cell-type across spots. We recommend trying both since fixed thresholding performs better for less common or cell-types which have low proportions everywhere while the otsu method performs better otherwise. Next, we use connected components analysis to group spots into regions [59]. All spots containing the celltype within a *D*_2−*nbd*_ *<*= *T*_2_ × *median*(*D*_2−*nbd*_) of other spots with the celltype were joined into a single region using breadth-first-search (BFS). We filter out regions containing 1 spot.

#### Cell subtypes identification in regions (CSI)

We transfer cell subtype annotations from the reference scRNA-seq to the spatial spots (Fig. 1h). In SpaDecoder, we learn a scRNA-seq cell by cell-type weight matrix Σ alongwith a cell-type proportion matrix *V* for each spot. We use Σ*V* for each spot, to derive a scRNA-seq cell by spatial spot mapping *M* for the entire dataset. We use this to map reference subtype annotations onto the spatial spots with *M* ^*T*^ *R*^*′*^ where *R*^*′*^ is the reference cell subtype annotation we wish to transfer. For each celltype region previously identified from CTRI, we assign a subtype label by comparing the proportions of each of the subtypes in a given spot under the assumption that the subtypes are not colocated in the same spot.

#### Predicting expression of Unseen Genes

In situations when all the genes are not captured spatially, we can use the cell to spot mapping *M* to predict the spatial expression patterns of all genes profiled by scRNA-seq (Fig 1i). Given the normalized reference scRNA-seq cell by gene expression matrix *B*^*′*^, we compute *Z* = *M* ^*T*^ *B*^*′*^ where *Z* is the spot by gene matrix *Z* containing all genes measured by scRNA-seq. These predicted expression profiles can further be normalized and used for downstream tasks such as finding spatial patterns of relevant markers and further for inferring cell-cell communication at a spot-neighborhood-celltype-pair-ligand-receptor resolution.

#### Identifying regions with varying cell type proportions across slices

Often, identifying regions of cell-type proportion changes across tissue slices is crucial to localizing developmental, disease-associated or tissue configuration induced spatial changes. To this end, we divided the spatial slice into regions and computed the Jenson-Shannon distance (JS) between the discrete probability distributions containing the celltype proportions for the same region across neighboring slices. We then permute the cell-type proportions (determined by SpaDecoder) between the spots within each slice and recompute the JS for each permutation to obtain a p-value.

#### Spatio-temporal location prediction

SpaDecoder can be applied to any stack of slices that can be well aligned. This further allows us to map reference scRNA-seq cells spatio-temporally (Fig. 1k). by mapping each scRNA-seq cell to the slice and spot with the highest mapping weight from the mapping matrix *M*.

### Data

To avoid tissue mismatch between reference scRNA-seq and query spatial datasets, we obtain matching scRNA-seq atlases, spatial datasets along with cell-type annotations from published sources. As discussed earlier, lack of ground truth cell type annotations in spots is a major limitation for evaluation of deconvolution algorithms. While ST simulation algorithms exist, spatial spots have varying number of cells and capturing the distance between spots, the mixture of cell-types in each spot, the cell-type density across the tissue, and spatial gene expression pattern of real ST tissues is challenging[60]. One solution is to simulate spots from real single-cell resolution ST assays by aggregating the expression profiles of cells in regions of varying sizes. We can then utilize the original published single cell resolution annotations to obtain spot level cell type proportions. A summary of the datasets used is in Supplementary Table 1. Preprocessing is performed in scanpy[31].

#### MERFISH of Mouse Hypothalamus Preoptic Region (*Moffitt2018*)

Spatial data was obtained from squidpy[57] via sq.datasets.merfish(). Ambiguous cell types and those missing from scRNA-seq data were excluded. For nomenclature consistency with scRNA-seq during evaluation, subtypes of endothelial (1, 2 and 3), immature oligodendrocyte (1, 2), and mature oligdendrocyte (3, 4) major cell types were grouped together into the corresponding major cell type. Blank genes were excluded.

scRNA-seq 10X reference data was downloaded from GSE113576 and associated metadata from Moffitt *et al* Table S1[35]. Cells having total counts *<* 2000 or *>* 25, 000, number of genes *<* 700 or mitochondrial counts *>* 10% were excluded. Doublets were separately detected for each sample with scrublet[61] and excluded. Cell types not present in any of the ST samples were excluded.

For visualization of the scRNA-seq UMAP embedding (Supp. Fig. 5a), cells were normalized with sc.pp.normalize_total, log1p transformed and batch corrected with combat using sc.pp.combat with each sample treated as a separate batch. Cell cycle phase scores and predicted phase for each cell were computed using scanpy.tl.score_genes_cell_cycle with lists of genes in S and G2M phases from (Table S2, Macosko *et al* [62]). Highly variable genes were obtained after excluding mitochondrial, ribosomal protein (Rpl and Rps), and mitochondrial ribosomal protein (Mrpl and Mrps) genes, using scanpy.pp.highly_variable_genes with default parameters. Principal component analysis (PCA) was performed with sc.tl.pca *svd_solver* = *arpack* and 40 PCs. The 10 nearest neighbor graph was used to perform partition-based graph abstraction (PAGA) which was used for initialization to obtain the UMAP embedding.

#### MERFISH of Mouse Retina (*Choi2023*)

The V45 integrated.h5ad file was downloaded from Zenodo[63] and separated into individual samples according to region and batch. 25 samples were selected as described in Supplementary Table 1. For the scRNA-seq reference, raw counts were obtained from GSE243413[38]. Similar to `Moffitt2018`, cells having total counts *<* 2000 or *>* 25, 000, number of genes *<* 700 or mitochondrial counts *>* 10% were excluded. We subset to reference cells with “GSE243413 (2023) WT_CD73” condition and visualized the remaining cells on the published embedding. The “majorclass” field was used for cell type annotation.

#### Xenium of Mouse model of Leptomeningeal metastasis (LM) melanoma (*Haviv2025*)

Raw scRNA-seq reference and spatial data were downloaded from Zenodo[64] with provided umap embedding and the “cell_type” annotation field.

#### Xenium of human breast cancer

The Xenium output bundle was downloaded from 10X[45]. zarr files for the 2 spatial samples were read with read_zarr function from thespatialdata[65] python package. The scRNA-seq reference h5 file alongwith associated annotations and the tSNE embedding for visualization. The “celltype” annotation field was used for both scRNA-seq and spatial data. Ambiguous cell types ‘Stromal_&_T_Cell_Hybrid’,’T_Cell_&_Tumor_Hybrid’ and those specific to a single modality were excluded. We merged fine grained cell type annotations and ran SpaDe-coder and other baselines on course annotations, leveraging fine grained scRNA-seq annotations to highlight our scRNA-spatial mapping and subtype annotation capabilities.

#### Visium of human fetal thymic development

The spatial thymus atlas was downloaded from cellxgene[47] and 16, 17, 18, 19 PCW samples were retained. scRNA-seq reference thymus atlas was also downloaded from cellxgene[66] and 15, 17, and 18 PCW samples were retained. The authors annotate cell types at various resolutions. We use “cell_type_level_0” annotations, excluding exploratory cell types annotated as “see_lv4_explore” but since we wish to study TEC compartments in greater detail, we choose “cell_type_level_2” annotations for Epithelial cells.

### Simulations

Spatial transcriptomic data inherently lacks ground truth cell-types in each spot. To this end, we simulate spots from single cell transcriptomic data for some experiments (Figs 2-4 and associated supplementary files). For Figure 2, we additionally simulate multiple slices from each single spatial sample to recapitulate the profiling of 3D tissue for evaluation purposes. The resulting raw spatial data was input to the preprocessing stage of SpaDecoder.

#### Spot simulation

We normalize the spatial coordinates to the range [0, 1]. Given our spot size parameter *N*_*spot*_, we divide each slice *i* from left right and top to bottom into square spots of side 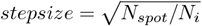 where *N*_*i*_ is the number of spots in slice *i*. Squares to the right and bottom are truncated to the spot size. Intuitively, *N*_*spot*_ is the expected number of cells in the area covered by a spot, assuming uniform distribution of cells throughout the tissue sample, so the square root gives us the side length. Since cells are not distributed uniformly we obtain spots containing different numbers of cells which recapitulates spatial sequencing technologies. Raw spot expression and spot coordinates are defined as the summed raw expression and averaged coordinates respectively of all cells in the spot. Ground truth cell type proportions are the average number of cells of each type in the spot. Default value is *N_spot* = 50. Experiments with other values are indicated.

#### Slice simulation

The first slice in the stack is chosen as the spatial sample. Each subsequent slice *i* is simulated by perturbing the previous slice *i* − 1. For each spot *j*, we identify the *k* + 1 nearest neighboring spots including the spot *j* itself. We randomly select *N*_*swap*_ cells in this neighborhood and randomly shuffle their indices such that their spot assignments and spatial locations are shuffled. The spot expression and cell type proportions are updated accordingly. Default number of neighbors *k* = 10 and *simulated slices* = 10. We performed experiments with varying *N*_*swap*_ values as described.

### Running SpaDecoder

All SpaDecoder experiments (except for testing parameter effects) are run with default parameters. Regularization parameter *λ* = 0.1. Adam optimization was performed with 500 iterations, early stopping if the cost function changes by *<* 10^−4^ between consecutive iterations and learning rate 0.01. In SpaDecoder-cluster (Supp. Fig 4), the cluster averaged version of SpaDecoder, each spot was initialized with cell type proportions from the reference. To prevent SpaDecoder from getting stuck in a local minima, deconvolution proportions for each spot were initialized with SpaDecoder-Cluster output. The scalar spot parameter *alpha* was initialized to 1. SpaDecoder was run in batch mode on an A40 GPU so multiple 3D spots could be processed in parallel. Permutation testing for the GearyC metric was also parallelized.

### Baseline Methods

#### CARD

CARD[11] was run in R 4.5.0 according to the [tutorial](https://yma-lab.github.io/CARD/documentation/04_CARD_Example.html). In particular, the steps included createCARDObject with minCountGene=1, minCountSpot=1 and no batch for consistent evaluation and CARD_deconvolution. The cell type proportions from Proportion_CARD were used for evaluation.

#### Cell2location

Cell2location[13] was run according to the [tutorial](https://cell2location.readthedocs.io/en/latest/notebooks/cell2location_tutorial.html) with default parameters. scRNA-seq reference preprocessing involved setting up the Regression Model with cell2location.models.RegressionModel.setup_anndata, RegressionModel, train, export posterior and obtaining the means_per_cluster_mu_fg. To run cell2location, the steps included cell2location.models.Cell2location.setup_anndata, cell2location.models.Cell2location, train() and export_posterior, with default parameters. The batch size was set to the total number of reference cells. “means_cell_abundance_w_sf” was extracted, normalized and used for evaluation.

#### Tangramsc and Tangram

Tangram and Tangramsc[6] were run according to the provided [tutorial](https://tangram-sc.readthedocs.io/en/latest/tutorial_link.html). Normalized spatial and scRNA-seq data was input to tg.pp_adatas. tg.map_cells_to_space was run on the GPU with the corresponding cluster key and mode=‘cells’ and ‘clusters’ for tangramsc and tangram respectively. tg.project_cell_annotations was run with the corresponding cluster key. Cell type predictions in ‘tangram_ct_pred’ were normalized and used for evaluation.

### Quantitative Evaluation

Quantitative evaluation was performed by comparing estimated cell type proportions from each of the methods with the cell types as annotated in the original publications with three metrics[21]: average root mean square error (RMSE), Jenson-Shannon divergence (JSD) and Pearson correlation.

#### Root Mean Square Error (RMSE)

For spot *j* spot in the *i*th slice, we compute Average 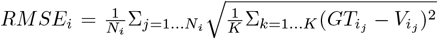 where *GT* corresponds to ground truth normalized cell type vector and *V* are those from the deconvolution method. *K* is the number of cell types, *N*_*i*_ the number of spots in the *i*th slice. Average RMSE is obtained by averaging per spot RMSE across all spots in the slice.

#### Jenson-Shannon divergence (JSD)

Let *D*_**p**,**q**_ = ∑_*k*_ (*p*_*k*_ ∗ *log*(*p*_*k*_*/q*_*k*_)) be the relative entropy between *p* and *q* where *p*_*k*_ and *q*_*k*_ are the respective probabilities. Let 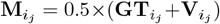 be the mean distribution between 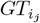 representing ground truth normalized proportion of cell types for spot *j* in slice *i* and 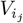 be the corresponding distribution from the deconvolution method. Then, for slice *i* with *N*_*i*_ spots, average 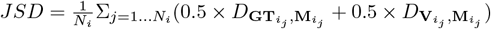.

#### Pearson correlation

Pandas corrwith with method=‘pearson’ was used to obtain Pearson correlation for each spot and averaged across all spots in each slice.

## Supplementary information

Supplementary Figures and Tables are attached with this manuscript.

## Acknowledgements

The authors thank the anonymous reviewers for their valuable suggestions.

## Declarations

## Data Availability

Publicly available data was used for this study. For `Moffitt2018`, spatial data was obtained from squidpy[57] via sets.merfish() and scRNA-seq 10X reference data was downloaded from GSE113576 with associated metadata from Moffitt et al Table S1[35]. For `Choi2023`, spatial data was downloaded from Zenodo[63] and scRNA-seq from GSE243413[38]. For `Haviv2025`, raw scRNA-seq reference and spatial data were downloaded from Zenodo[64]. For the human breast cancer dataset[45], the Xenium output bundle was downloaded from 10X. For the human thymic development dataset, the spatial thymus atlas was downloaded from cellxgene[47] and scRNA-seq reference from cellxgene[66].

## Code Availability

The code for the SpaDecoder package and applications is available at https://github.com/ZhangLabGT/spadecoder.

## Author Contribution

XZ, ZZ and ML conceived the idea, ML and ZZ implemented the package, ML performed tests and drafted the manuscript. XZ and ZZ supervised the project. All authors revised and approved the manuscript.

