## Supplementary Information for "Spatiotemporal cell type deconvolution leveraging tissue structure"

#### Supplementary Note 1

A summary of the notation used in SpaDecoder is included here. $G$: number of common selected genes in the reference scRNAseq and spatial dataset

$\boldsymbol{G_{sc}}$: set of genes in reference scRNAseq dataset $P$: number of slices in the stack

$N_i$: number of spots in $i$th slice in stack

$\boldsymbol{X_i}$: $G \times N_i$ gene expression matrix for $i$th slice in stack

$\boldsymbol{x_{i_j}}$: $G$ length gene expression column vector for $j$ spot in $i$th slice of stack

Hence, $\boldsymbol{X_i} = [\boldsymbol{x_{i_1}}, \boldsymbol{x_{i_2}}, ..., \boldsymbol{x_{i_{N_i}}}]$

$\boldsymbol{X} = (\boldsymbol{X_1}, \boldsymbol{X_2}, ..., \boldsymbol{X_P})$: set containing expression matrix stack for $P$ slices ordered according to tissue z-axis

$M$: number of cells in reference scRNAseq dataset

$K$: number of celltypes in reference scRNAseq dataset

$\boldsymbol{B_{sc}}$: $G \times M$ matrix of expression of cells in reference scRNAseq

$\boldsymbol{B_{bulk}}$: $G \times K$ matrix of per celltype average expression of cells in reference scRNAseq

$\boldsymbol{B_{avg}}$: $G \times K$ matrix of per celltype average expression $\boldsymbol{C}$: $M \times K$ one-hot cell-type label matrix defined as

$$\mathbf{C}_{st} = \begin{cases} 1 & \text{if cell } s \text{ is of type } t \\ 0 & \text{otherwise} \end{cases} \tag{1}$$

$\boldsymbol{\Sigma_{i_j}}$: **learned** $M \times K$ cell type label association matrix for spot $j$ in slice $i$

$\alpha_{i_j}$: **learned** scalar for spot $j$ in slice $i$

$\boldsymbol{\alpha_i} = [\alpha_{i_1}, \alpha_{i_2}, ..., \alpha_{i_{N_i}}]^T$: $N_i$ length column vector with $\alpha_{i_j}$'s for $N_i$ spots in slice $i$

$\boldsymbol{\alpha} = (\boldsymbol{\alpha_1}, \boldsymbol{\alpha_2}, ..., \boldsymbol{\alpha_P})$: set containing stack of $\boldsymbol{\alpha_i}$'s

$\boldsymbol{v_{i_j}}$: $K$ length **learned** cell type proportion column vector for $j$th spot in $i$th slice

$\boldsymbol{V_i} = [\boldsymbol{v_{i_1}}, \boldsymbol{v_{i_2}}, ..., \boldsymbol{v_{i_{N_i}}}]$: $K \times N_i$ matrix of cell type proportions in $N_i$ spots of slice $i$

$\boldsymbol{V} = (\boldsymbol{V_1}, \boldsymbol{V_2}, ..., \boldsymbol{V_P})$: set containing stack of $\boldsymbol{V_i}$'s

$\lambda_{i_j}$: regularization parameter for spot $j$ in slice $i$

$\boldsymbol{a_{i_j}}$: length $M$ scRNAseq cell mapping column vector for spot $j$ in slice $i$

$\boldsymbol{D_{gene}} = \{scRNAseq\ cell\ associated\ gene\ features\}$. Set of gene features in scRNAseq data. For example $d \in \boldsymbol{D_{gene}}$ maybe a gene captured in the scRNAseq dataset

$\boldsymbol{D_m} = \{scRNAseq\ cell\ associated\ metadata\}$. Set of metadata features in scRNAseq data. For example $d \in \boldsymbol{D_m}$ maybe a cell type or subtype annotation

$\boldsymbol{r_d}$: length $M$ scRNAseq cell vector for some feature $d \in (\boldsymbol{D_{gene}} \cap \boldsymbol{D_m})$

$\boldsymbol{s}$: $M$ length size factor vector from total count normalization of scRNAseq cells

### Supplementary Figures and Tables

**Supplementary Table 1** Overview of datasets used in this study

| Ref. | Species | Ref. Samples | Cells | Spatial Dataset | Spatial Samples | Genes |
|---|---|---|---|---|---|---|
| [1] | mouse | all | 25326 | Merfish hypothalamus[1] | 12 | 135 |
| [2] | mouse | GSE243413 WT CD73 | 20155 | Merfish mouse retina[3] | 10, 14-16, 20-26, 40-46, 50-56 (Total:25) | 500 |
| [4] | mouse | all | 9855 | Xenium mouse LM metastasis[4] | 1 | 243 |
| [5] | human | all | 26031 | Xenium breast cancer[5] | 2 | 313 |
| [6] | human | 15, 17, 18 PCW | 34798 | Visium thymic development[7] | 16-19 PCW | 36406 |

**Supplementary Table 2** Summary of simulations and figures

| Dataset | Simulation | Parameters/Datasets | Slices | Figure |
|---|---|---|---|---|
| Moffitt2018[1] | Spots+slices | $N_{spot} = 50$; $N_{swap} = 2$; stack=10 | 132 | Fig. 2, Supp Fig. 1-6, 8-10 |
| Moffitt2018[1] | Spots+slices | $N_{spot} = 50$; $N_{swap} = 5$; stack=10 | 132 | Fig. 2, Supp Fig. 1-2, 5, 8 |
| Moffitt2018[1] | Spots+slices | $N_{spot} = 50$; $N_{swap} = 10, 20$; stack=10 | 132 | Supp. Fig. 7-8 |
| Moffitt2018[1] | Spots+slices | $N_{spot} = 50$; $N_{swap} = 50$; stack=10 | 132 | Supp. Fig. 8 |
| Moffitt2018[1] | Spots+slices | $N_{spot} = 10, 25, 50, 75, 100$; $N_{swap} = 2$; stack=10 | 132 | Supp. Fig. 9 |
| Choi2023[2, 3] | Spots+slices | $N_{spot} = 50$; $N_{swap} = 2$; stack=10 | 275 | Fig. 2, Supp Fig 1-6, 8-10 |
| Choi2023[2, 3] | Spots+slices | $N_{spot} = 50$; $N_{swap} = 5$; stack=10 | 275 | Fig. 2, Supp Fig 1-2, 5, 8 |
| Choi2023[2, 3] | Spots+slices | $N_{spot} = 50$; $N_{swap} = 10, 20$; stack=10 | 275 | Supp Fig. 7-8 |
| Choi2023[2, 3] | Spots+slices | $N_{spot} = 50$; $N_{swap} = 50$; stack=10 | 275 | Supp Fig. 8 |
| Choi2023[2, 3] | Spots+slices | $N_{spot} = 10, 25, 50, 75, 100$; $N_{swap} = 2$; stack=10 | 275 | Supp Fig. 9 |
| Haviv2025[4] | Spots+slices | $N_{spot} = 50$; $N_{swap} = 2$; stack=10 | 11 | Fig. 2, Supp Fig. 1-6, 8-10 |
| Haviv2025[4] | Spots+slices | $N_{spot} = 50$; $N_{swap} = 5$; stack=10 | 11 | Fig. 2, Supp Fig. 1-2, 5, 8 |
| Haviv2025[4] | Spots+slices | $N_{spot} = 50$; $N_{swap} = 10, 20$; stack=10 | 11 | Supp. Fig. 7-8 |
| Haviv2025[4] | Spots+slices | $N_{spot} = 50$; $N_{swap} = 50$; stack=10 | 11 | Supp. Fig. 8 |
| Haviv2025[4] | Spots+slices | $N_{spot} = 10, 25, 75, 100$; $N_{swap} = 2$; stack=10 | 11 | Supp. Fig. 9 |
| Moffitt2018[1] | Spots | $N_{spot} = 50$ | 12 | Fig. 3, Supp. Fig. 11 |
| Janesick2023[5] | Spots | $N_{spot} = 50$ | 2 | Fig. 4, Supp. Fig. 12 |
| Yayon2024[7] | NA | NA | 4 | Fig. 5, Supp. Fig. 13 |

**a**

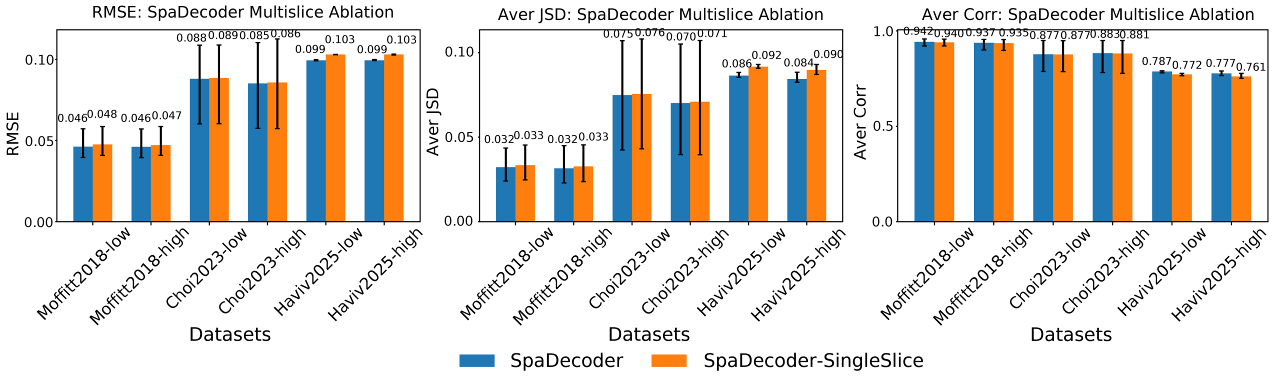


**Supplementary Figure 1 Utilizing multiple slices improves deconvolution - SpaDecoder multislice ablation in support of Fig. 1** (a) Barplots showing the average RMSE, JSD and Pearson correlation of SpaDecoder and SpaDecoder-SingleSlice with simulated spots (spot size parameter $N_{spot} = 50$) and simulated slice stacks $N_{swap} = 2$ (suffix -low) and $N_{swap} = 5$ (suffix -high) on Moffitt2018[1], Choi2023[3], and Haviv2025[4]. SpaDecoder-SingleSlice follows SpaDecoder except that it only uses the current slice for deconvolution

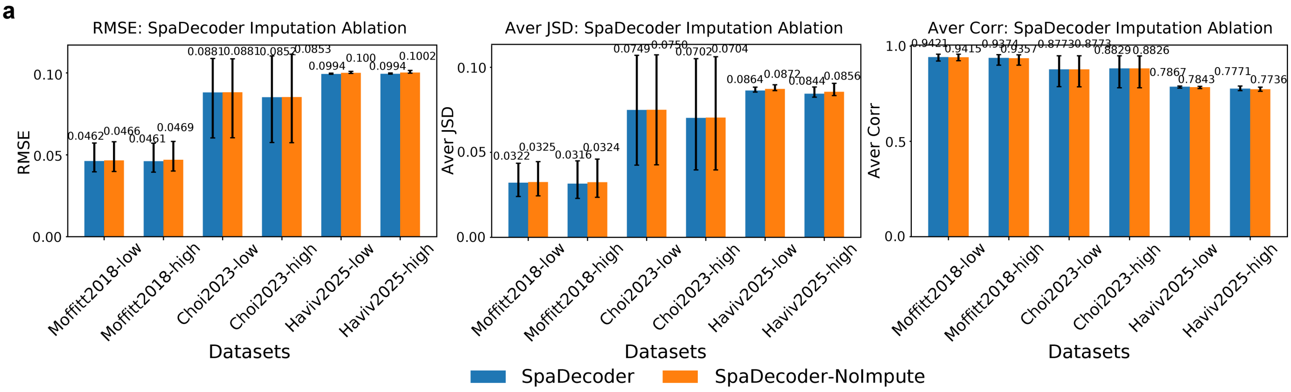


**Supplementary Figure 2 Imputing intermediate slices improves deconvolution - SpaDecoder imputation ablation in support of Fig. 1** (a) Barplots showing the average RMSE, JSD and Pearson correlation of SpaDecoder and SpaDecoder-NoImpute with simulated spots (spot size parameter $N_{spot} = 50$) and simulated slice stacks $N_{swap} = 2$ (suffix -low) and $N_{swap} = 5$ (suffix -high) on Moffitt2018[1], Choi2023[3], and Haviv2025[4]. SpaDecoder-NoImpute follows SpaDecoder except that it does not use intermediate imputed slices for deconvolution

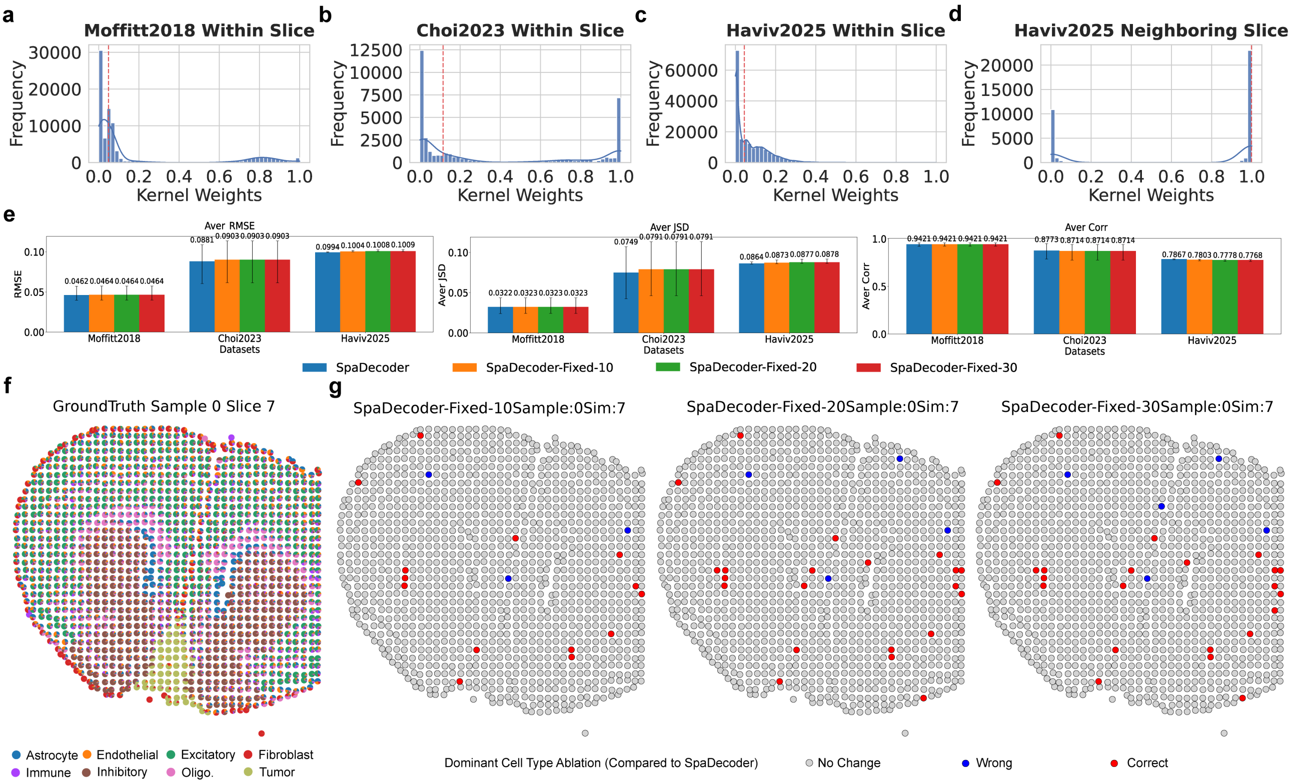


**Supplementary Figure 3 Variable 3D Spatio-transcriptomic kernel improves performance - SpaDecoder adaptive kernel ablation, in support of Fig. 1** (a-d) Distribution of within-slice kernel weights for Moffitt2018 (a), Choi2023 (b), Haviv2025 (c) and neighboring slice kernel weights for Haviv2025 (d) (e) Barplots showing the average RMSE, JSD and Pearson correlation of SpaDecoder and fixed spatial neighbor ablations with 10 (SpaDecoder-Fixed-10), 20 (SpaDecoder-Fixed-20) and 30 (SpaDecoder-Fixed-30) on Moffitt2018[1], Choi2023[3], and Haviv2025[4]. $N_{spot} = 50$, $N_{swap} = 2$. (f-g) Spatial visualization of representative sample 0, slice 7 in stack colored by pie plot visualizations of cell type proportions in each spot obtained from original annotations (ground truth) (f) and mismatches in the dominant cell-type detected by SpaDecoder and the ablations in (e). Gray indicates alignment in both SpaDecoder and the ablation, blue shows a wrong type identified by the SpaDecoder and red is a mistake in the ablation correctly identified by SpaDecoder (g)

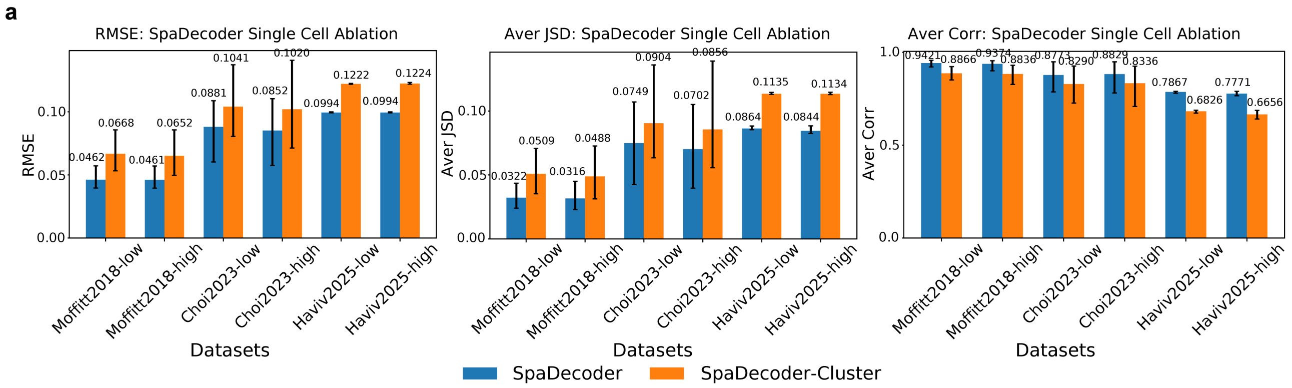


**Supplementary Figure 4 Single cell resolution scRNAseq reference improves deconvolution - SpaDecoder single cell reference ablation in support of Fig. 1** (a) Barplots showing the average RMSE, JSD and Pearson correlation of SpaDecoder and SpaDecoder-Cluster with simulated spots (spot size parameter $N_{spot} = 50$) and simulated slice stacks $N_{swap} = 2$ (suffix -low) and $N_{swap} = 5$ (suffix -high) on Moffitt2018[1], Choi2023[3], and Haviv2025[4]. SpaDecoder-Cluster follows SpaDecoder except that it uses cluster averaged scRNAseq reference expression profiles instead of single cell resolution

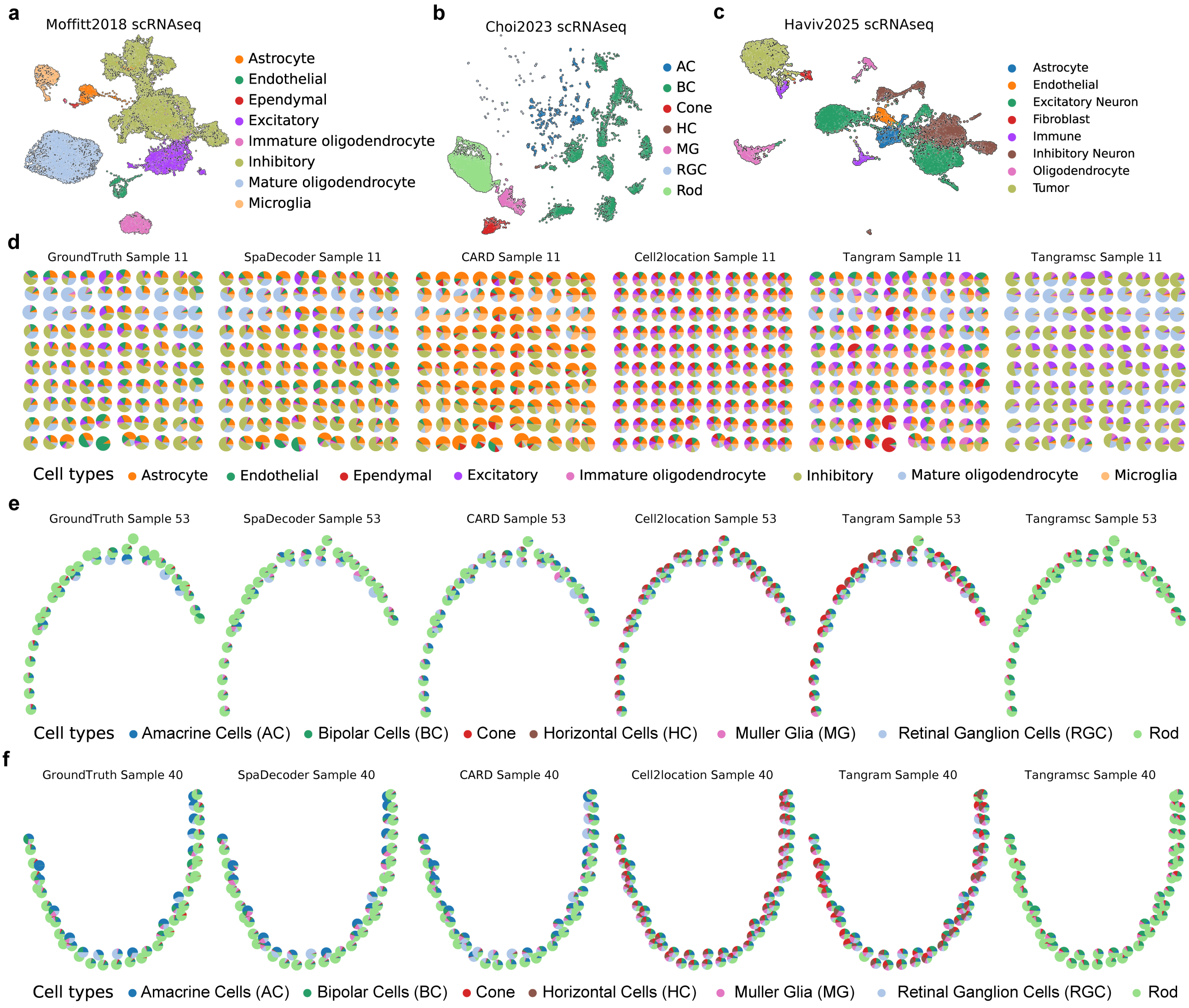


**Supplementary Figure 5 (in support of Fig. 2)** (a-c) UMAP embedding of scRNAseq reference cells colored by annotations from Moffitt et al (Moffitt2018)[1] (a) Choi et al (Choi2023)[3] (b) and Haviv et al (Haviv2025)[4] (c) (d) Pie plot visualizations of cell type proportions in each spot obtained from original annotations (ground truth) in Moffitt2018[1], SpaDecoder, CARD, Cell2location, Tangram, and Tangramsc for representative sample 11 (e-f) Pie plot visualizations of cell type proportions in each spot obtained from original annotations (ground truth) in Choi2023[3], SpaDecoder, CARD, Cell2location, Tangram, and Tangramsc for representative sample 53 (e) and 40 (f)

a

b

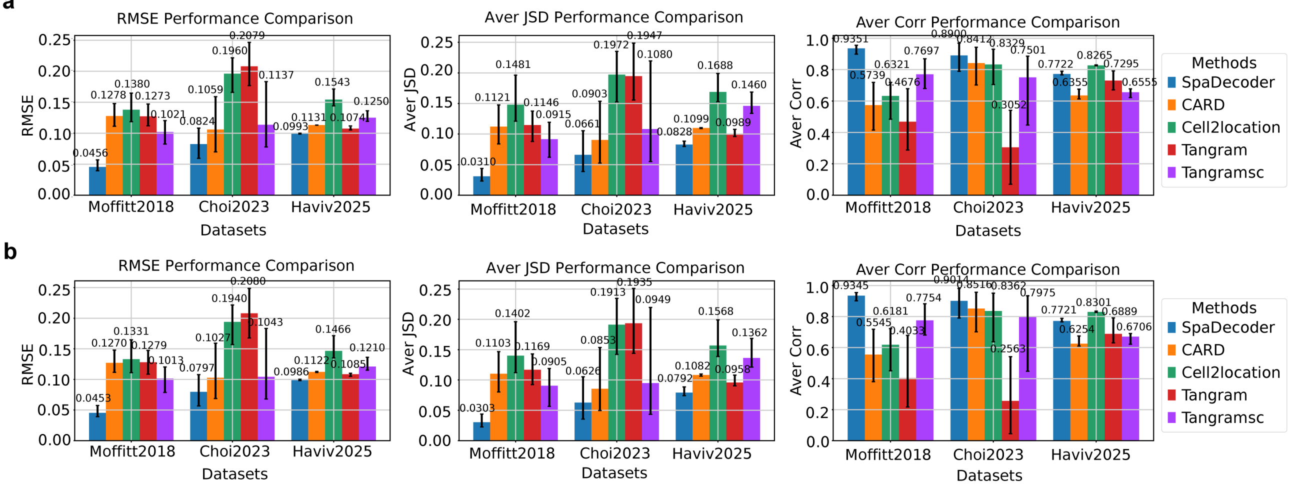


**Supplementary Figure 6 (in support of Fig. 2) Performance evaluation with various slice stack simulations** (a-b) Barplots showing the average RMSE, JSD, Pearson correlation of SpaDecoder, CARD, Cell2location, cluster-averaged Tangram and single cell resolution Tangram (Tangramsc) across 3 single cell spatial datasets Moffitt2018[1], Choi2023[3], and Haviv2025[4] with simulated spots (spot size parameter $N_{spot} = 50$) and simulated slice stacks $N_{swap} = 10$ (a) and $N_{swap} = 20$ (b)

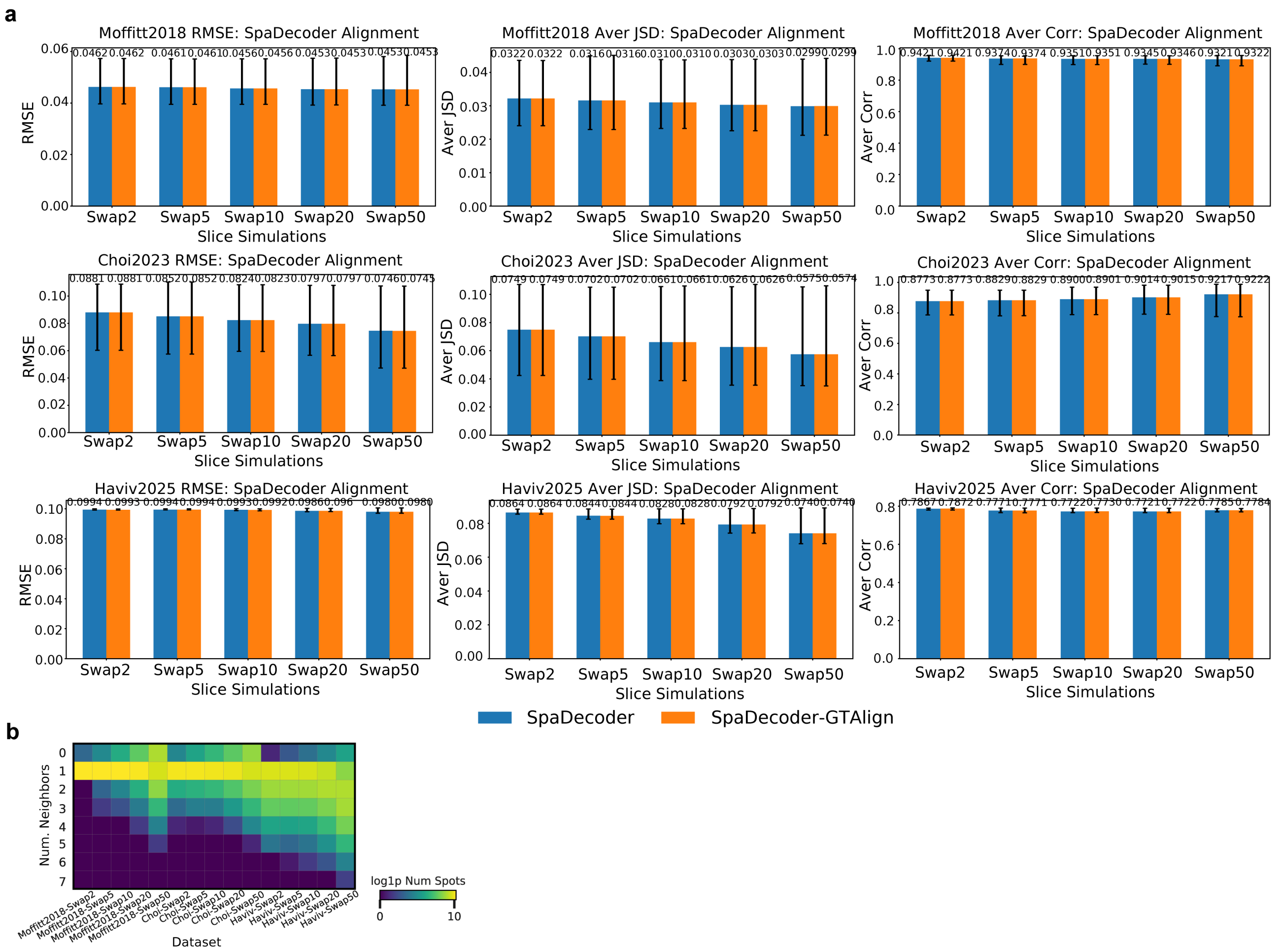


**Supplementary Figure 7 (in support of Fig. 2) Demonstrating the utility of alignment and adjacent slice neighborhood selection** (a) Barplots showing the average RMSE, JSD and Pearson correlation of SpaDecoder and SpaDecoder-GTAlign with simulated spots (spot size parameter $N_{spot} = 50$) and simulated slice stacks $N_{swap} = 2, 5, 10, 20, 50$ on Moffitt2018[1], Choi2023[3], and Haviv2025[4]. SpaDecoder-GTAlign follows SpaDecoder except that it aligns corresponding ground truth spots in the slice pair instead of inferring alignment with moscot[8] (b) Heatmap showing the $log(1 + number\ of\ central\ spots)$ having a selected adjacent neighborhood of $y$ spots (along Y-axis) for a given slice pair across the various datasets and simulation conditions (along X-axis).

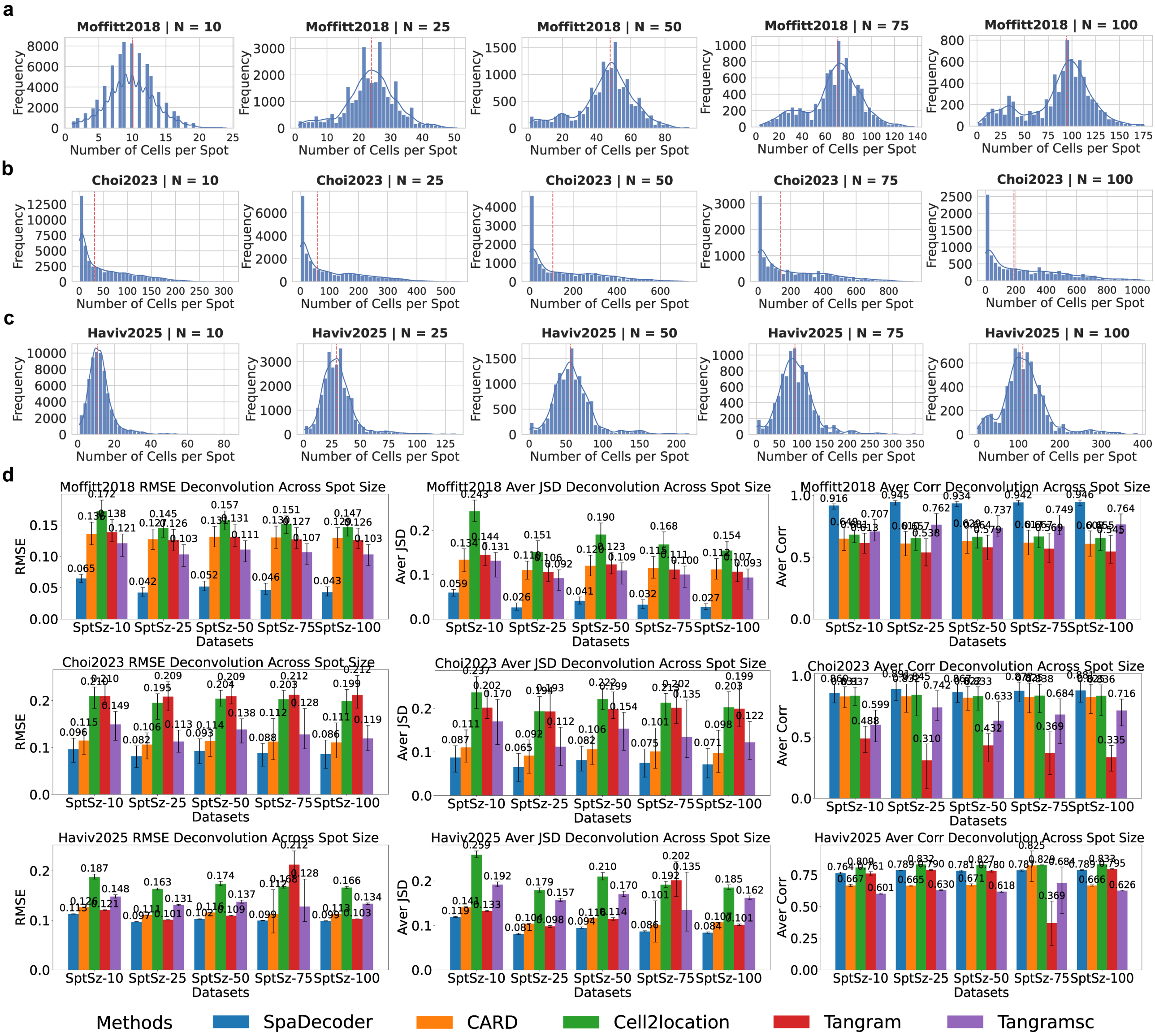


**Supplementary Figure 8 (in support of Fig. 2) Robustness of SpaDecoder on benchmarks with varying spot size across datasets** (a) Histogram showing the number of cells in each spot across simulation conditions with spot parameter $N_{spot} = 10$, $N_{spot} = 25$, $N_{spot} = 50$, $N_{spot} = 75$, $N_{spot} = 100$ on Moffitt2018[1], Choi2023[3], and Haviv2025[4]. (b) Barplots showing the average RMSE, JSD and Pearson correlation of SpaDecoder, CARD, Cell2location, Tangram, and Tangramsc with simulated spots using spot size parameters from (a)

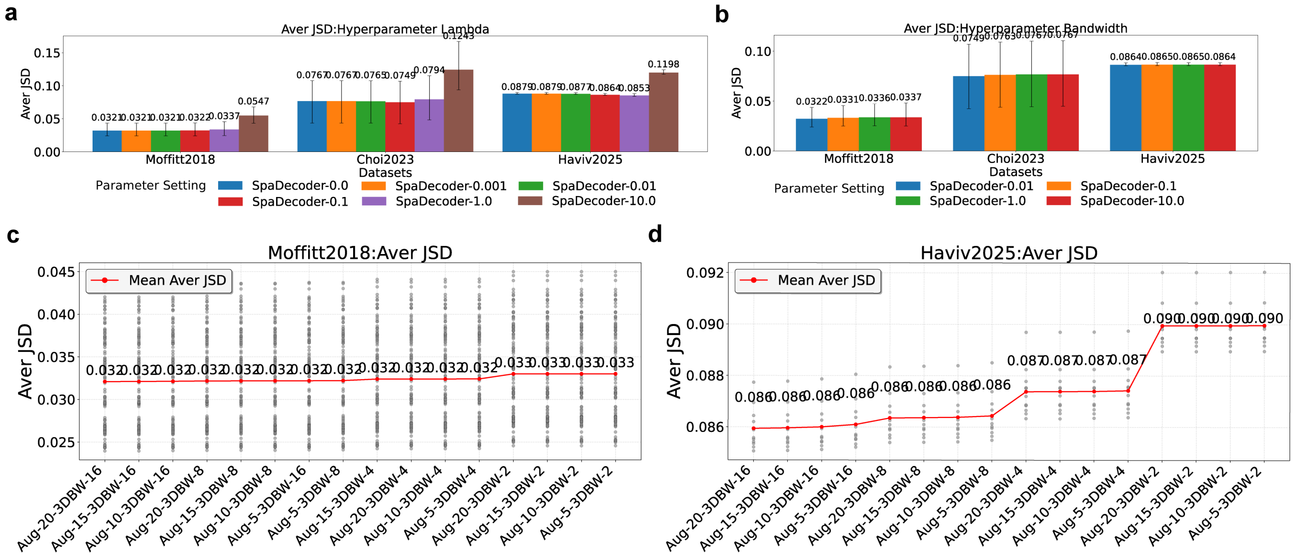

**Supplementary Figure 9 (in support of Fig. 2) Spadecoder is stable to ranges of hyperparameter choices** (a-b) Barplots showing the average JSD for SpaDecoder deconvolution on Moffitt2018[1], Choi2023[3], and Haviv2025[4] with multiple hyperparameter choices for $\lambda$ (a) and 3D bandwidth (b). (c-d) Plots showing the average JSD for SpaDecoder deconvolution on Moffitt2018[1] (c) and Haviv2025[4] (d) obtained by varying the hyperparameters for number of augmented slices between every slice pair (Aug-) and 3D kernel bandwidth (3DBW-). Each point in the plot corresponds to the average JSD on a single slice in the 3D stack. Mean average JSD trends across the parameter choices are visualized in red.

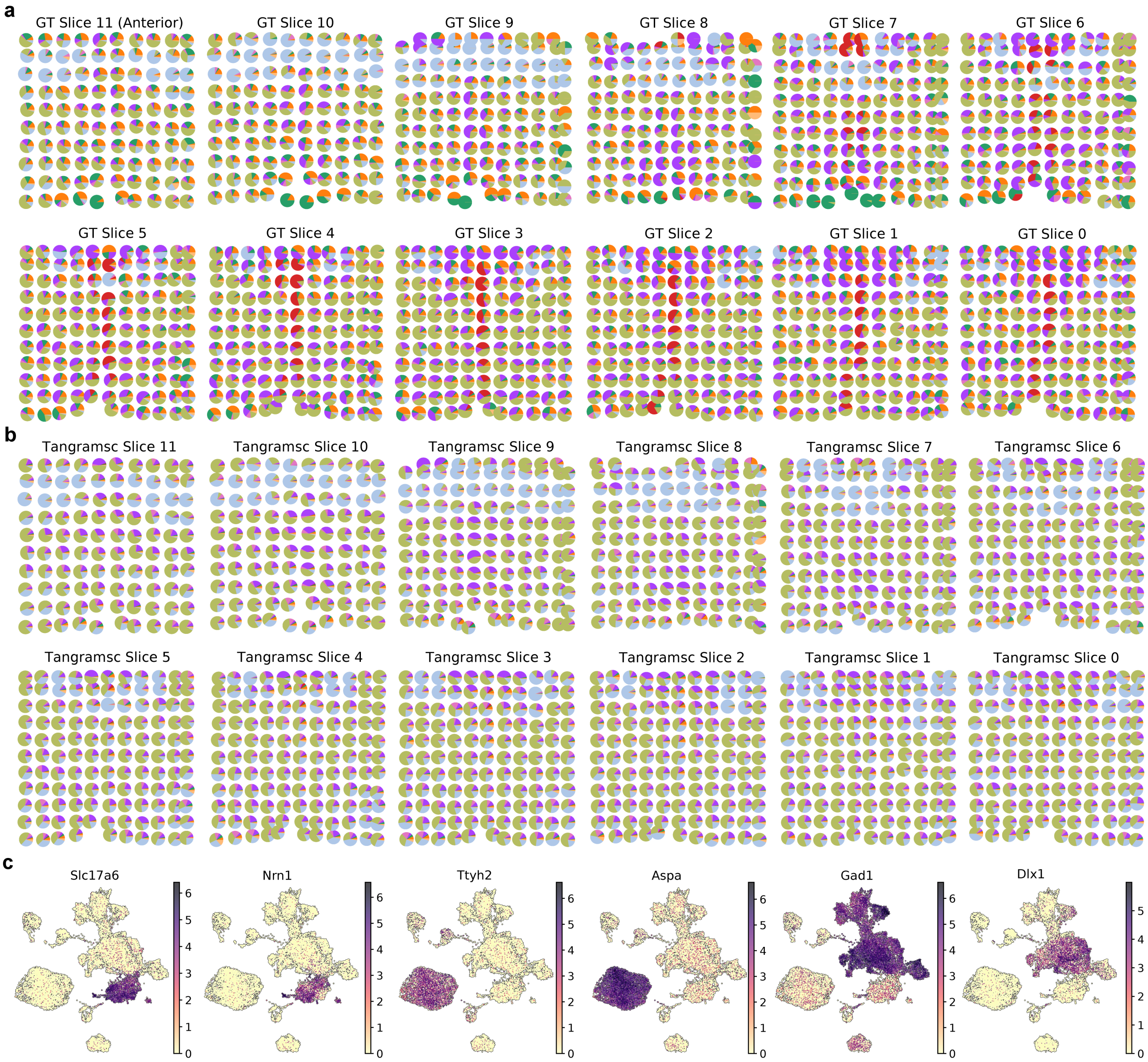


**Supplementary Figure 10 (in support of Fig. 3)** (a-b) Pie plot visualizations of cell type proportions obtained from original annotations (ground truth) in Moffitt2018 (a) and Tangramsc estimated cell type proportions (b) for all 12 slices along the anterior (slice 11) to posterior (slice 0) axis (c) UMAP of Moffitt2018 scRNAseq reference with cells colored by log2-normalized expression values of key literature driven celltype specific markers for Excitatory neurons (*Slc17a6*, *Nrn1*), Mature Oligodendrocytes (*Ttyh2*, *Aspa*) and Inhibitory neurons (*Gad1*, *Dlx1*)

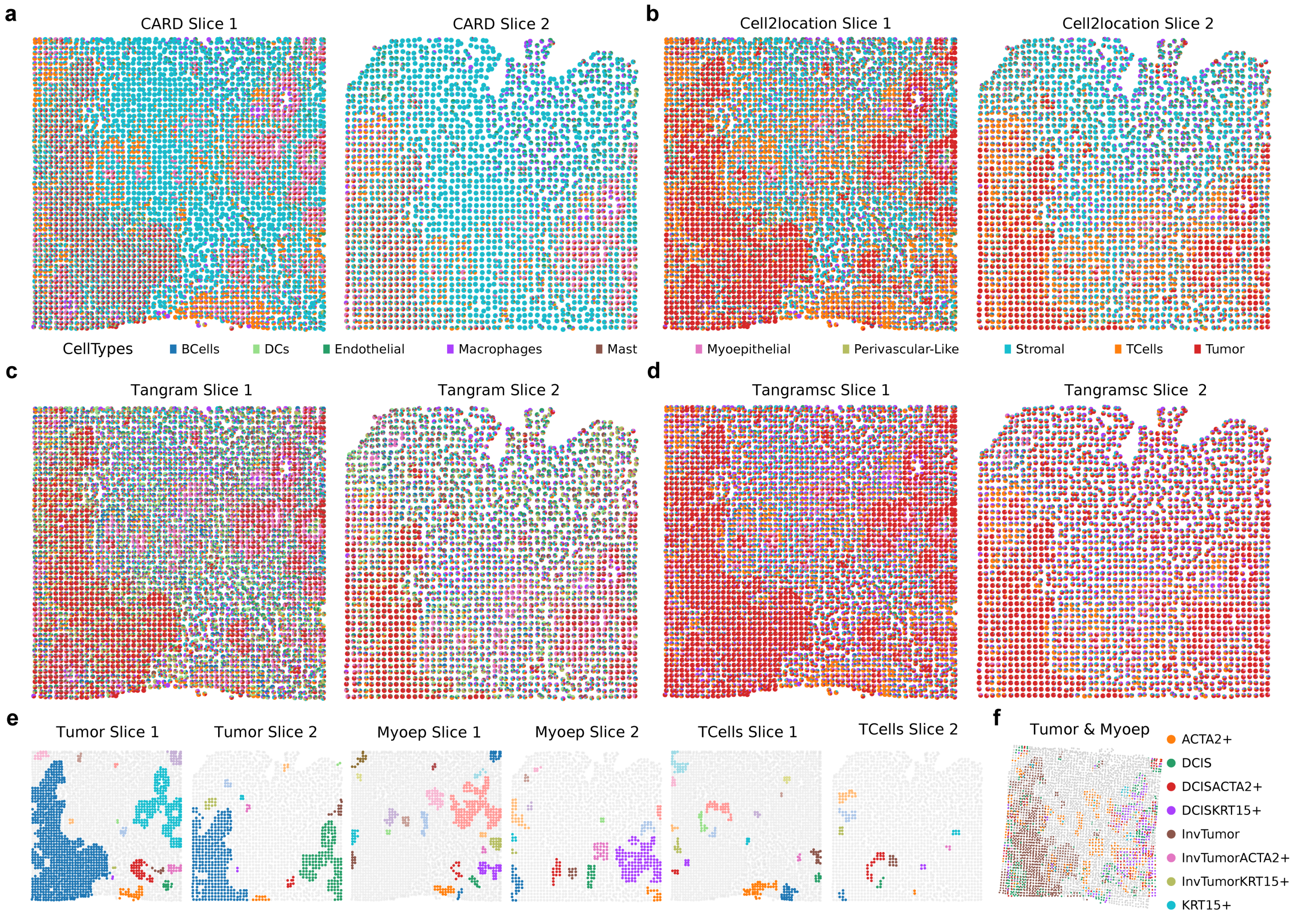


**Supplementary Figure 11 (in support of Fig. 4)** (a-d) Pie plots of spot cell type proportions from CARD (a), Cell2location (b), Tangram (c) and Tangramsc (d) on two Xenium slices from Janesick et al[5] with spots obtained from aggregating cells in neighborhoods (e) Spatial slices colored by tumor, ME, and Tcell regions obtained from connected component analysis of corresponding SpaDecoder celltype proportions (f) Aligned spatial slices colored by Tumor/ME subtype from identified regions. Spots containing both tumor and ME subtypes are labeled accordingly.

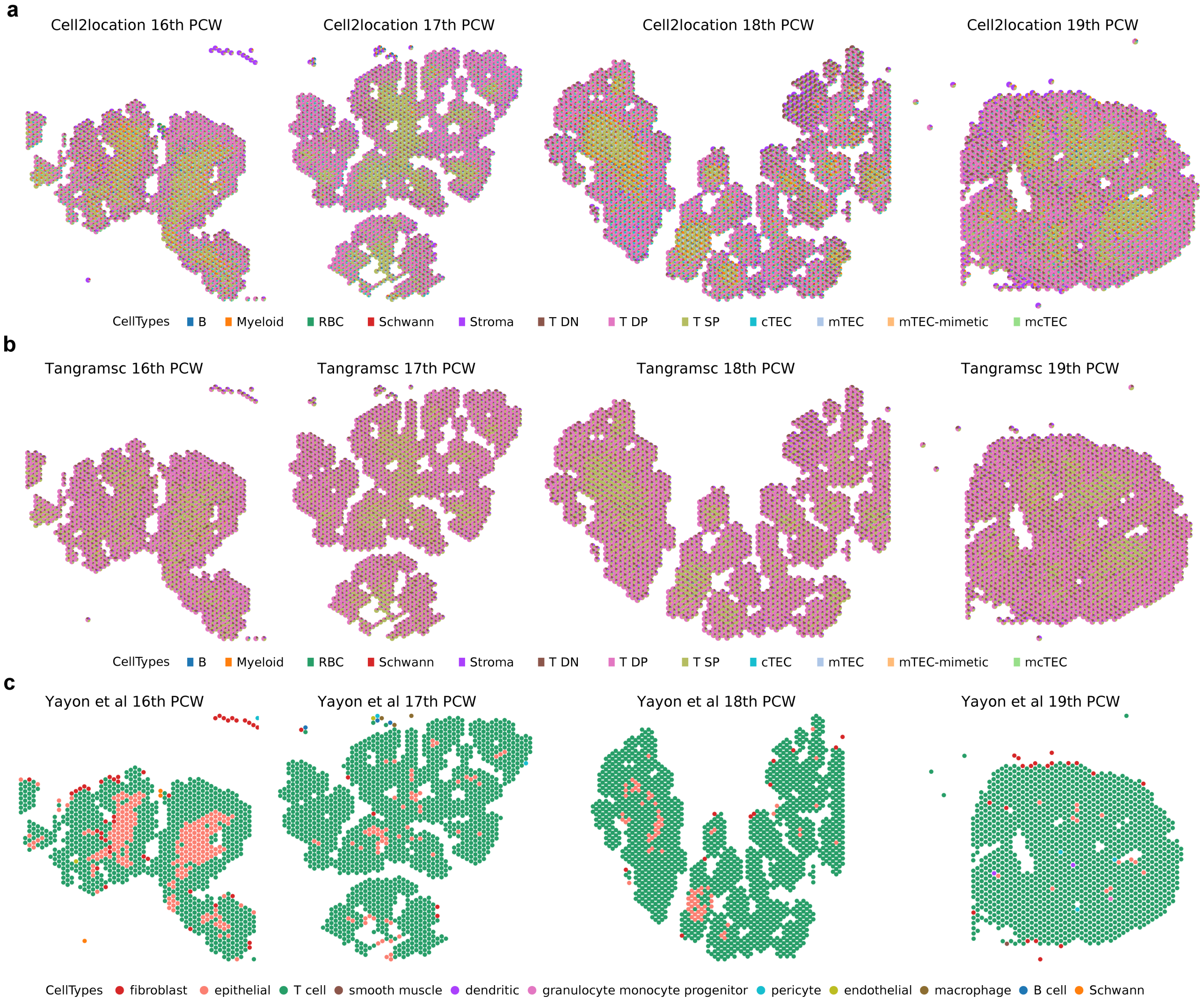


**Supplementary Figure 12 (in support of Fig. 5)** (a-b) Pie plots of cell type proportions on Visium slices from 16-19PCW of human fetal thymic development estimated by Cell2location (a) and Tangramsc (b) (c) Spatial plots of human fetal thymic development from 16 to 19PCW indicating cell types as annotated by Yayon et al[7] with each spot colored by a single cell type
